# Dominant and Genome-Wide Formation of DNA:RNA Hybrid G-Quadruplexes in Living Yeast Cells

**DOI:** 10.1101/2023.02.16.528764

**Authors:** Chen-xia Ren, Rui-fang Duan, Jia Wang, Yu-hua Hao, Zheng Tan

## Abstract

Guanine-rich nucleic acids form G-quadruplex (G4) structures that play a critical role in cellular processes. Previous studies have mostly focused on monomeric intramolecular G4s with four consecutive guanine tracts (G-tracts) from a single strand. However, this structural form has never been confirmed in eukaryotic cells. Here, we report the formation of hybrid G4s (hG4s), consisting of G-tracts from both DNA and RNA, in the genome of living yeast cells. Analysis of Okazaki fragment syntheses and G4-specific probing reveals that hG4s can efficiently form with as few as a single DNA guanine-guanine (GG) tract due to the participation of G-tracts from RNA. This finding increases the number of G4-forming sites in the yeast genome from 38 to 587,694, a more than 15,000-fold increase. Interestingly, hG4s still form and even dominate at genomic G4 sites that are theoretically capable of forming the monomeric intramolecular DNA G4s by themselves. Compared to DNA G4s (dG4s), hG4s exhibit a wider range of kinetics, higher prevalence, and greater structural diversity and stability. Most importantly, hG4 formation is tightly coupled to transcription through the involvement of RNA, allowing hG4s to function in a transcription-dependent manner. Overall, our study establishes hG4s as the overwhelmingly dominant G4 species in the yeast genome and emphasizes a renewal of the current perception of the structural form, formation mechanism, prevalence, and functional role of G4s in eukaryotic genomes. It also provides a sensitive and currently the only method for detecting the structural form of G4s in living cells.

**Significance:** The identification of hybrid G-quadruplexes (hG4s) has disclosed a previously unrecognized structural form of G4s as the most common and abundant G4 species in the yeast genome. It reveals not only a dominant rule governing the formation of G4s in eukaryotic genomes, but also a unique genotype that allows G4-mediated transcriptional regulation to take feedback from the output as input, thus allowing the creation of feedback loops at the transcriptome scale.

## Introduction

Guanine-rich nucleic acids have the ability to form four-stranded G-quadruplex (G4) structures in which four guanine tracts (G-tracts) are bundled by stacked guanine tetrads (G-tetrads) (Figure 1A). Due to their diverse topology, physical stability, and specific genomic location, G4s play critical roles in various cellular processes, including replication, transcription, translation, and genome stability (1,2). Putative G-quadruplex sequences (PQSs) are found throughout the genomes of both prokaryotic and eukaryotic organisms (3). Importantly, PQSs are not randomly distributed but are highly concentrated in regulatory regions, particularly in promoters of higher organisms, suggesting their involvement in the regulation of gene expression (4,5). Currently, our understanding of G4s in living cells is mainly extrapolated from *in vitro* studies. However, G4 formation in cells occurs in a completely different environment, which has been shown to cause G4s to behave differently in terms of kinetics, conformation, stability, and other properties than they do under simplistic *in vitro* conditions (6).

**Figure 1.**
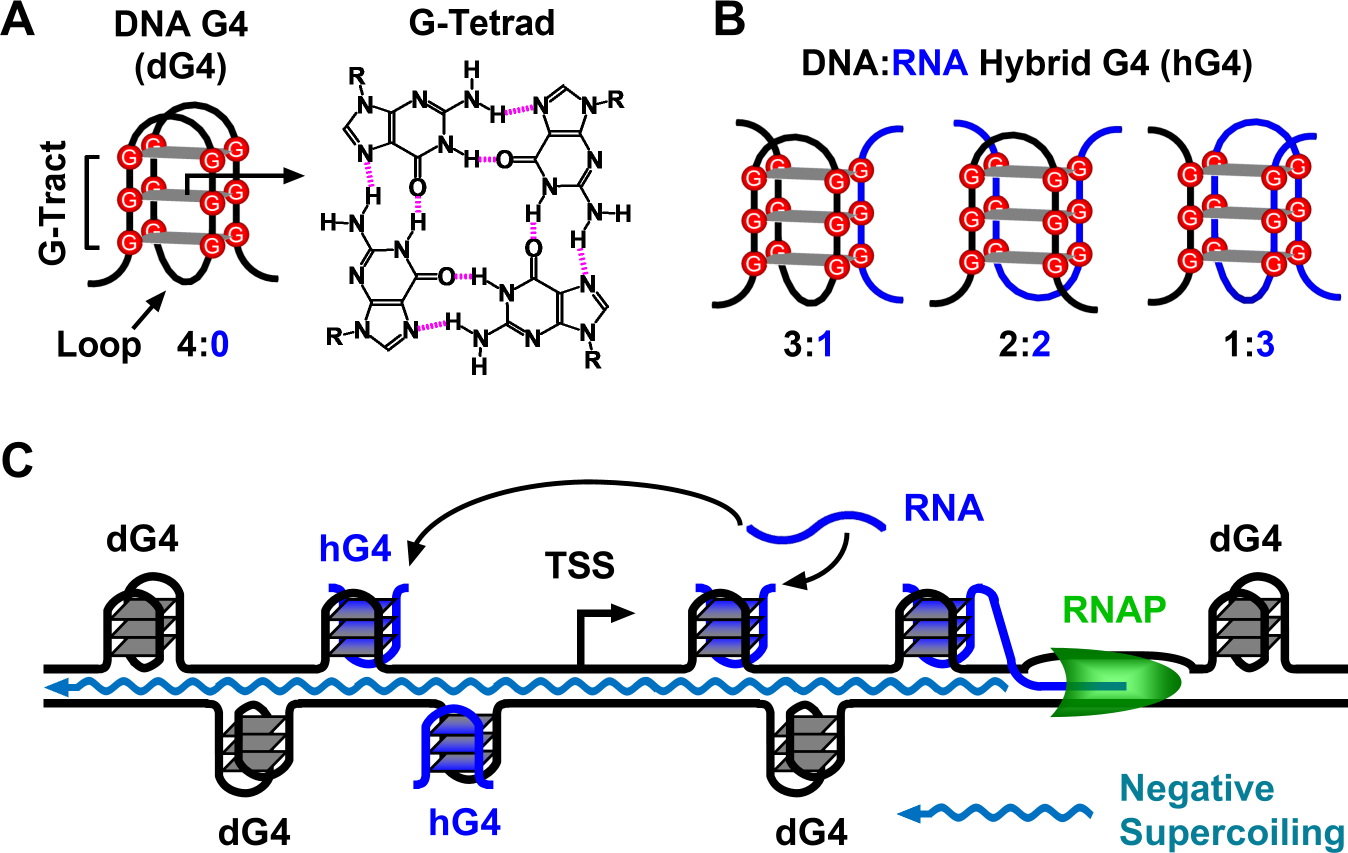
Intramolecular and intermolecular G4 structures and their formation in a DNA duplex transcribed *in vitro*. (A) Intramolecular G4 composed of four G-tracts with three G-tetrads. (B) DNA:RNA hybrid G4s can be formed in transcribed DNA by the joint participation of G-tracts from DNA and RNA with different strand stoichiometry. (C) G4 formation can be triggered by a transcribing RNA polymerase (RNAP) on its upstream side through the propagation of negative supercoiling and on its downstream side by an approaching transcription bubble. See text for further details.

In principle, the four G-tracts of a G4 can originate from a single or multiple nucleic acid strands, resulting in either an intramolecular or intermolecular structure. Research in G4 biology has primarily focused on canonical intramolecular DNA or RNA G4s, where four G-tracts are derived from a single strand (Figure 1A), as highlighted in recent reviews (4,5). Although G4s have recently been detected in the genomes of living animal cells (7), their actual form has not been determined. A decade ago, our *in vitro* studies demonstrated the formation of a specific type of G4s, termed DNA:RNA hybrid G4s (hG4s) (Figure 1, B-C), in transcribed DNA. This was achieved by the simultaneous recruitment of G-tracts from a non-template DNA strand and RNA transcripts (8,9). In our previous studies (3,8–16), we characterized hG4s in terms of their formation mechanism, kinetics, stability, structural diversity, competition with DNA G4s (dG4s), and other features. Interestingly, it was found that hG4s can still form and even dominate in DNAs with four G-tracts, although these DNAs are theoretically capable of forming intramolecular dG4s by themselves (12). This phenomenon was later explained by *in vitro* studies showing that hG4s are more stable and fold faster than the corresponding dG4s (16–20). Although we have confirmed the formation of hG4s in living bacterial cells (12), it is still unclear whether such structures can be formed in eukaryotic cells.

The recruitment of G-tracts from RNA allows DNA to form a G4 with as few as one DNA G-tract, instead of four (Figure 1B). This results in a significantly higher number of hG4 sites in a genome compared to dG4 sites (9). For example, in the yeast genome, the number of hG4 PQSs is over 3,300 times greater than that of dG4 PQSs, considering only those capable of forming a G4 with three or more G-tetrads. Previous *in vitro* studies have shown that hG4 formation is directly proportional to transcriptional activity (13,21), indicating a tight coupling between hG4s and transcription. Therefore, hG4 formation represents a unique genotype that is significantly more abundant and likely plays a distinct role in conjunction with transcription. However, uncertainty about its existence has prevented researchers from exploring and understanding this unique genotype in the eukaryotic kingdoms. The answer to the question of whether hG4 can form in eukaryotic cells should reveal an *in vivo* reality, as well as a unique regulatory mechanism that has gone virtually unnoticed. In this study, we present evidence for and characterize the genome-wide formation of hG4s in the yeast genome and discuss its implications for genome-related cellular activities.

## Materials and Methods

### Polymerase and exonuclease stop assay

The two assays were conducted using Bsu DNA polymerase, large fragment (NEB, USA) and Exo I exonuclease (NEB, USA) with the DNAs listed in Table S1, as previously described (22,23).

### Data from public repositories

The yeast chromosome sequences (sacCer3) and bed files for yeast SGD genes were downloaded from the UCSC Genome Browser (http://genome-asia.ucsc.edu). Five paired-end OK-Seq datasets (GSE115897, GSE118078, GSE139860, GSE141884, GSE173065) of Okazaki fragments (OKFs) in either bed or bedpe format (24–28) were downloaded from the Gene Expression Omnibus database (GEO, https://www.ncbi.nlm.nih.gov/geo/) hosted on the National Center for Biotechnology Information (NCBI) website. Data in bedpe format were converted to bed format using a custom-modified Perl script that we adapted from bedpe2bed.pl, which is available at https://github.com/srinivasramachandran/sam2bed/blob/main/bedpe2bed.pl.

### Identification of PQS and control motifs in genome

A regular expression, G{3,}({1,7}?G{3,})}0,}, was used to search the chromosome sequences for PQSs containing one or more G_≥3_ tracts as described (3). The number of G-tracts in the PQSs found was determined by the regular expression G{3,4}. For example, a G_7_ PQS would be identified as having two G_3_ tracts. Based on such analyses, we organized the resulting PQSs into multiple bed files based on the number of G-tracts. Each bed file was then divided into two separate bed files; one containing PQSs on the forward strand and the other containing PQSs on the reverse strand.

A similar approach was used to identify PQSs containing one or more G_≥2_ tracts with the regular expression G{2,}(.{1,7}?G{2,}){0,}. Subsequently, the PQSs identified were searched with the regular expression G{3,} to remove motifs containing any G_≥3_ tract, leaving only those that could only form hG4s or dG4s of only two G-tetrads. DNA motifs containing one or more A_≥3_ tracts were identified using the regular expression A{3,}(.{1,7}?A{3,}){0,}. Motifs containing one or more GNG tracts, where N is any non-G nucleotide, were identified using the regular expression [^G]{7}G[^G]G(.{1,7}? G[^G]G){0,} [^G]{7}.

### Identification of Orphan PQSs

PQS bed files were processed by the Bedtools merge software (29) with the arguments “-d n -c 4,5,6 -o count,collapse,distinct”, where the “-d n” represents the maximum distance “n” between PQSs to be merged. PQSs that were not merged and flagged with a count of one were collected as “orphan PQSs”.

### Survey of PQSs in genes

The gene bed files were extended by 120 bp on the upstream side to include the promoters (30,31) using the Bedtools slop software (29). The number of PQSs overlapping with the extended gene regions by at least one bp was counted using the Bedtools map software (29).

### Distribution of Okazaki fragments (OKFs) across PQSs

The bed files of OKFs were converted to bedgraph format using the Bedtools genomeCoverageBed software (29) to represent the full length (-bg -strand +/-), 3’ (-bg −3-strand +/-), or 5’ (-bg −5 -strand +/-) ends of the OKFs, respectively. The resulting bedgraph files were then converted to bigwig format using the Bedtools bedGraphToBigWig software (29) with a bin size of 10 nucleotides (nts), unless specified otherwise. The distribution of OKFs and their 3’ or 5’ ends over the 3’-end of PQSs was profiled using the computeMatrix and plotProfile tools of the Deeptools software (32), along with the corresponding bigwig and PQS bed files. The final profile for each dataset was an average of all samples.

To profile the 5’ end of the OKFs whose 3’-end was within ±20 nts of the 3’-end of a PQS, a new PQS bed file was created to represent the PQS 3’ ±20 nts regions. A new OKF bed file was also created by copying columns 2 and 3 to columns 7 and 8 and then modifying columns 2 and 3 to represent the OKF 3’-ends. The overlap of regions between the two bed files was determined using the intersect command of the Bedtools software (28) to remove the OKFs in the OKF bed file whose 3’-end did not overlap with any regions in the PQS bed file. Columns 7 and 8 of the OKF bed file were then restored to columns 2 and 3, respectively. The resulting OKF bed file was then processed to profile the distribution of the 5’- and 3’-ends of the OKFs at the PQS 3’-ends as described in the previous section.

### Plasmid construction for G4 probe (G4P) ChIP

The coding sequence of the G4P protein was amplified from pNLS-G4P-IRES2-EGFP (7) using a PCR primer pair of 5’-ATTAAGCTTATGCCCAAGAAGAAGCGGAAG-3’and 5’-GTGGATCCTTACTTGTCATCGTCATCCTT-3’and inserted into the pYES2 plasmid between the Hind III and BamH I sites.

### G4P ChIP-Seq

The recombinant plasmid pYES2-G4P-3xFLAG was introduced into *S. cerevisiae* strain BY4741 using the lithium acetate/PEG transformation method (33). The transformant was inoculated on SC-Ura agar medium (synthetic dropout agar medium without uracil). Colonies were collected after 72 hrs of incubation at 30 °C and then cultured in SD-Ura liquid medium (synthetic complete medium without uracil supplemented with 2% (w/v) glucose) at 30 °C to an OD_600_ of 2. Cells were then harvested by centrifugation and washed three times in SG-Ura liquid medium (synthetic complete medium without uracil supplemented with 2% (w/v) galactose). Cell pellets were resuspended in liquid SG-Ura to an OD_600_ of 0.6 and incubated at 30 °C to activate the GAL1 promoter.

After 6 hrs of G4P induction, cells were cross-linked with 1% formaldehyde and then mixed with 425-600 µm glass beads (Cat# G8080, Solarbio, China) and lysed using a bead ruptor (Bioprep-6, Allsheng, Hangzhou, China) at 6.0 m/s for 8 cycles of 30 sec on at 4 µC and 10 min off on ice. The resulting lysate was pelleted, resuspended in 600 µl of lysis buffer and then sheared using a Covaris M220 ultrasonicator for 20 min at 4-7 °C (10% duty cycle, 75 W power, 200 burst).

For DNA sequencing, 1% of the DNA fragment sample was saved as input, and the remainder was immunoprecipitated using 30 µl anti-FLAG M2 magnetic beads (Cat# M8823, Sigma-Aldrich) according to the protocol (34). Purified DNA was sequenced on an Illumina HiSeq platform (Genewiz, Suzhou, China) to generate 2×150 bp paired-end reads for input and G4P-bound DNA. Clean fastq data were then aligned to the UCSC sacCer3 genome using the Bowtie2 software and mapped to PQSs or other indicated motifs as described (7).

For sequencing RNA in hG4s, 5 µg of monoclonal DNA antibody (Cat# MA1-83116; Invitrogen) was incubated for 2 hrs at 4 °C with Protein A/G Mix Magnetic Beads (Cat# LSKMAGAG10, Millipore). After washing with dilution buffer, the beads were mixed with DNA fragments and immunoprecipitation (IP) was performed as described (34). The resulting elution, without cross-linking reversal, was split into two halves, one half incubated with 20 µl mouse IgG anti-FLAG M2 magnetic beads for G4 pull-down and the other half incubated with 20 µl mouse IgG magnetic beads (Cat#5873; Cell Signaling Technology) for non-specific control. Immunoprecipitation was conducted as described (34). The elution was subjected to cross-linking reversal and then treated with DNase I and Proteinase K, respectively. The RNA was purified using an RNA MinElute column (Cat# 74204; Qiagen) and was further treated with DNase I. The remaining RNA was then purified using the RNA MinElute column. RNA library was prepared and strand-specific RNA-Seq was performed on an Illumina HiSeq platform (Genewiz, Suzhou, China).

To determine the presence of PQSs in hG4 RNAs, the R1 reads of the RNA-Seq data in fastq format were first processed using the fastp software with the following options: “--length_required 5 --length_limit 55 --dedup”, with the remaining options left as defaults. The resulting fastq data was then aligned to the UCSC sacCer3 genome using the Hisat2 software with the options “-k 1 --fast --no-softclip --no-unal”. The samtools view software was then used with the options “-q 20 -F 0×100” to filter for the aligned RNA reads, discarding unaligned reads and ensuring that each read was only reported once, if it could be aligned to multiple positions in the genome. The resulting bam files were converted to fasta format using the Samtools view and awk commands. For reads aligned in the reverse direction, their sequences were reverse-complemented using the Seqkit seq software. Finally, the PQS motifs in the fasta files were identified as described previously (9).

### Distribution of G4P across PQSs

Distribution of G4P binding across regions of interest was calculated as described (7).

### Brief background on G4 formation *in vitro*

Our group has performed most of the studies (3,8–16) on hG4s (3,8–20), including their discovery, identification, mechanism of formation, and effect on transcription. The co-transcriptional formation of G4s is a dynamic process that can be driven by different mechanisms. On the non-template strand downstream of a moving RNA polymerase (RNAP), G4 formation is triggered when an approaching transcription bubble is 7 nucleotides (nts) away from a PQS (35) (Figure 1C). Alternatively, a G4 can form behind a moving RNAP driven by a negative supercoiling wave (21,36), which can propagate upward and induce G4 formation over a range of several thousand nts (36) (Figure 1C). In the case of hG4s, they can begin to form during the second round of transcription when RNA produced on the template strand in the previous round is displaced to provide G-tracts (10). In addition, an hG4 can form on the upstream side of transcription with G-tracts in RNA produced by transcription on the downstream side (14). Compared to dG4s, hG4s have a faster folding rate and greater mechanical stability (16), allowing hG4s to dominate even when a DNA itself is capable of forming a dG4 (12). Taken together, these observations suggest that hG4s can be formed regardless of the genomic location of DNA G-tracts, as long as there is an adequate supply of RNA with G-tracts from the cellular RNA pool.

## Results

### G4 detection based on stop of Okazaki fragment (OKF) synthesis

Protein translocation along a DNA strand is impeded by a G4 structure (37), which has led to the development and widespread use of a polymerase stop assay (22) for the *in vitro* detection of G4s in both DNA (22,38,39) and RNA (40,41). In this method, a short DNA primer is annealed to a target strand and extended by DNA polymerase. The extension stops upon encountering a G4 to signal the presence of a G4 structure. As shown in Figure 2 (A and B), a yeast or human DNA G4 halted the progress of both DNA polymerase and exonuclease. In contrast, both reactions reached their full length when the PQSs were mutated to prevent G4 formation. In Figure 2C and 2D, the DNA template and substrate each contained seven consecutive G-tracts, allowing the formation of only one dG4 at four alternative positions (42). Consequently, four DNA bands corresponding to the dG4 at the four alternative positions are observed in both assays. This feature has been used to identify the G-tracts involved in G4 formation (12,23).

**Figure 2.**
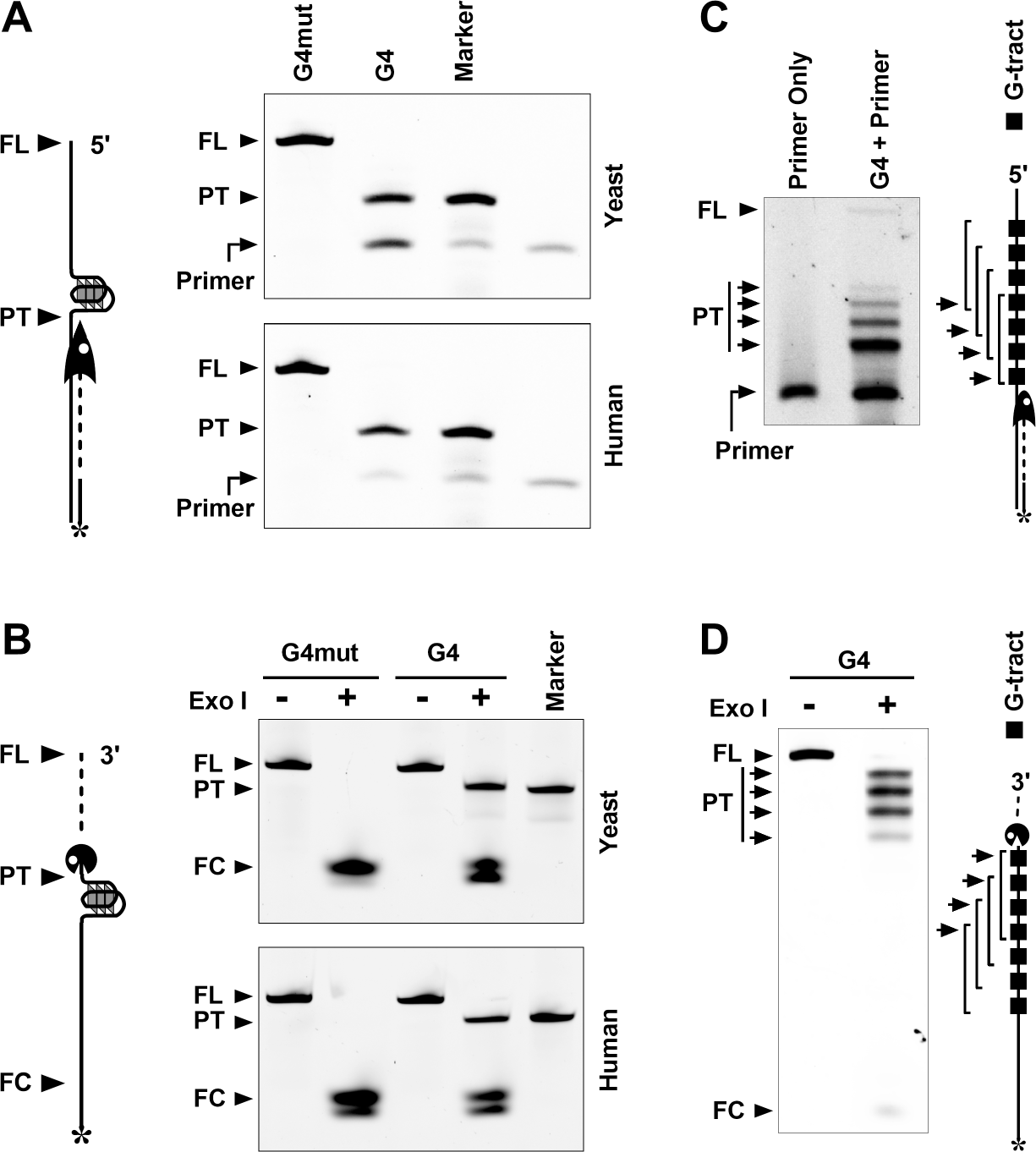
Stop of DNA polymerase and exonuclease by an intramolecular dG4. (A) DNA polymerase stop assay showing premature termination (PT) of DNA synthesis caused by a G4 and full-length (FL) extension with a G4 mutant. (B) DNA exonuclease stop assay showing premature termination (PT) of DNA cleavage caused by a G4 and full-length cleavage (FC) with a G4 mutant. The G4 PQS was derived from the promoter of the human c-KIT gene and the region between the yeast ATO3 and EFT2 genes, respectively. (C) DNA polymerase stop assay showing premature termination (PT) of DNA synthesis caused by a G4 that can form at multiple alternative positions (brackets). (D) DNA exonuclease stop assay showing premature termination (PT) of DNA cleavage caused by a G4 that can form at multiple alternative positions (brackets).

To detect G4 formation in living yeast cells, we used the *in vitro* polymerase stop principle to monitor the progress of Okizaki fragment (OKF) synthesis primed by RNA instead of DNA. OKFs are short DNA fragments generated within a replication fork on the lagging strand during DNA replication (43). In recent years, OKF synthesis in yeast has been extensively studied using the OK-Seq technique on other topics in several independent investigations (24–28), which provided us with the original OK-Seq data for this study.

### Retardation of OKF synthesis by PQSs

A stable G4 usually has three or more G-tetrads. We first identified PQSs in the yeast genome and classified them into four groups, 4G3+, 3G3+, 2G3+, and 1G3+, based on the number of G_≥3_ tracts. While the 4G3+ PQSs can form dG4s by themselves, the 3G3+, 2G3+, and 1G3+ PQSs have the potential to form hG4s by recruiting additional G-tracts from RNA, as shown in Figure 1B. We then profiled the distribution of OKFs relative to the 3’-ends of each PQS group (Figure 3A) using OK-Seq data from the GEO database. Regardless of the G4 type that the PQSs could form, all profiles showed a positive peak downstream of the PQSs on the PQS-bearing strands (Figure 3, B-E, red lines). These positive peaks showed an abrupt decrease at the 3’-end of the PQSs, suggesting that OKF synthesis was intercepted at the 3’ front of the dG4s in the 4G3+ PQSs (Figure 3B) and hG4s in the 3G3+, 2G3+, and 1G3+ PQSs (Figure 3, C-E), respectively, as in the *in vitro* polymerase stop assay (Figure 2) (22).

**Figure 3.**
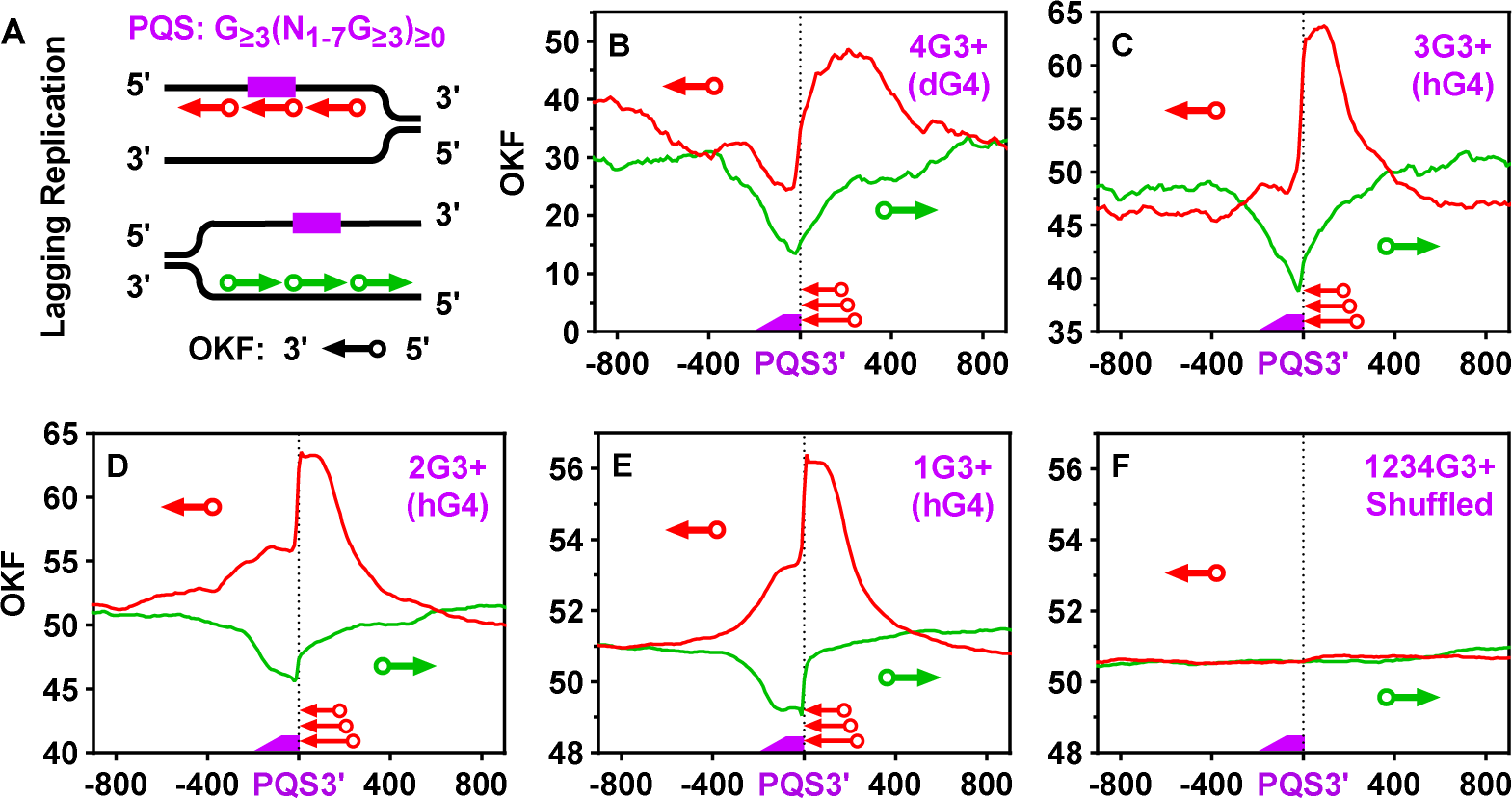
(A) Scheme of Okazaki fragment (OKF) distribution profiling on the PQS-bearing strand and the strand opposite to a PQS. (B-E) Distribution of OKFs at the 3’-end of PQSs with 1 to 4 or more G_≥3_ tracts and capable of forming (B) dG4s or (C-E) hG4s of three or more G-tetrads. (F) Distribution of the 3’-end of OKFs at the 3’-end of randomly shuffled PQSs. Original data for this and OKF-related figures are from GSE115897 (24) unless otherwise noted. The “3+” sign indicates that the G-tracts in the PQS had three or more consecutive guanines; the digit before the “G” indicates the number of G-tracts in the PQS; text in parentheses indicates the type of G4 the PQS can form. Bin size: 10 nts.

In contrast to the PQSs, the complementary cytosine-rich (C-rich) motifs instead were associated with a negative OKF peak (Figure 3, B-E, green lines), because they do not form G4s, while their flanking regions had scattered background G4 formation. The two oppositely polarized OKF peaks were specifically associated with PQSs, as they both disappeared when the coordinates of the PQSs were randomly shuffled across the genome and the OKF signal was re-profiled over the resulting fake PQSs (Figure 3F). Similar profiles were obtained with four additional independent datasets from the GEO database (Figure S1-S4), suggesting that the retardation of OKF synthesis at the 3’-end of PQSs is a universal phenomenon.

To further verify the G-tract dependence, we performed a similar analysis using motifs in which the guanines were substituted with adenines. In this case, a negative peak was observed on both DNA strands for the OKF (Figure S5). This result was expected since the A/T-rich motifs lacked the ability to form a G4, while their flanking regions had a scattered background of G4 formation. Introducing a mutation in the middle of a G-tract effectively disrupts G4 formation (Figure 2). We also examined motifs containing one or more GNG tracts, where N represents any nucleotide other than G. Since these motifs were also unable to form G4, they also exhibited a negative OKF peak on both DNA strands (Figure S6). As expected, shuffling these two types of motifs resulted in the disappearance of the negative peaks (Figure S5F and Figure S6F). Taken together, these results support the formation of hG4s in the motifs containing G-tracts (Figure 3, Figure S1-S4).

### Confirmation of G4 formation with a G4 probe (G4P) protein

The G-tract-dependent arrest of OKF synthesis at the PQSs provides evidence for the formation of G4 structures in these regions (22). To confirm this, we expressed a small G4P protein in yeast cells and performed G4P ChIP-Seq to examine G4 formation (Figure 4), following a similar approach used in human and other animal cells (7). The G4P, with its two G4-binding domains and a molecular weight of only 6.7 kDa (Figure 4A), exhibits high specificity for different G4s. Consistent with the OKF signal (Figure 3), the G4P showed enrichment at the PQSs, which could form either dG4s (Figure 4B) or hG4s (Figure 4, C-E). This enrichment disappeared when the coordinates of the PQSs were randomly shuffled, and the G4P signal was re-profiled (Figure 4F). We also analyzed the G4P signal over the A/T-rich and GNG motifs. Consistent with the OKF signals at these motifs (Figure S5 and Figure S6), G4P also showed a negative peak (Figure S7 and Figure S8), further confirming the formation of hG4 and dG4 in the PQSs.

**Figure 4.**
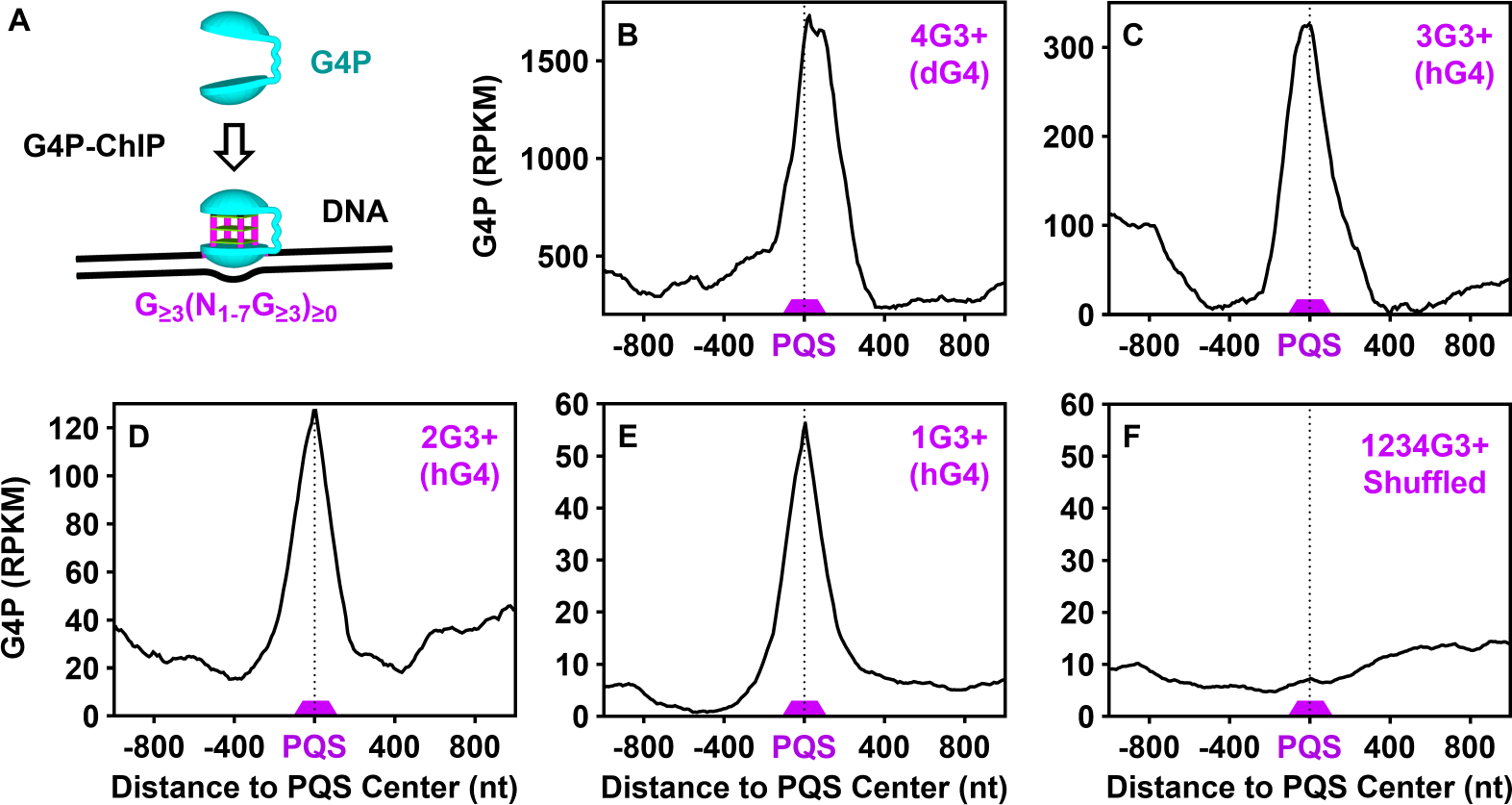
(A) Scheme of G4 detection by G4P ChIP-Seq. (B-E) Enrichment of G4P at PQSs with 1 to 4 or more G_≥3_ tracts and capable of forming (B) dG4s or (C-E) hG4s of three or more G-tetrads. (F) Distribution of G4P at randomly shuffled PQSs. PQSs are the same as in Figure 3.

### Formation of hG4s and dG4s of two G-tetrad layers

While the G4 community has primarily focused on stable G4 structures with three or more G-tetrads, our previous study revealed the formation of hG4s with only two G-tetrads in DNA duplexes transcribed *in vitro* (8). These hG4s are less stable than those with three or more G-tetrads (8,13). We were intrigued to investigate whether such hG4s could also form in living cells. Therefore, we analyzed OKF and G4P signals around PQSs containing only GG tracts. The results showed an accumulation of OKF (Figure 5) and an enrichment of G4P (Figure 6) at such PQSs, providing evidence for the formation of these hG4s and dG4s in the genome. Based on this discovery, our subsequent analysis of G4 will include those with two G-tetrads.

**Figure 5.**
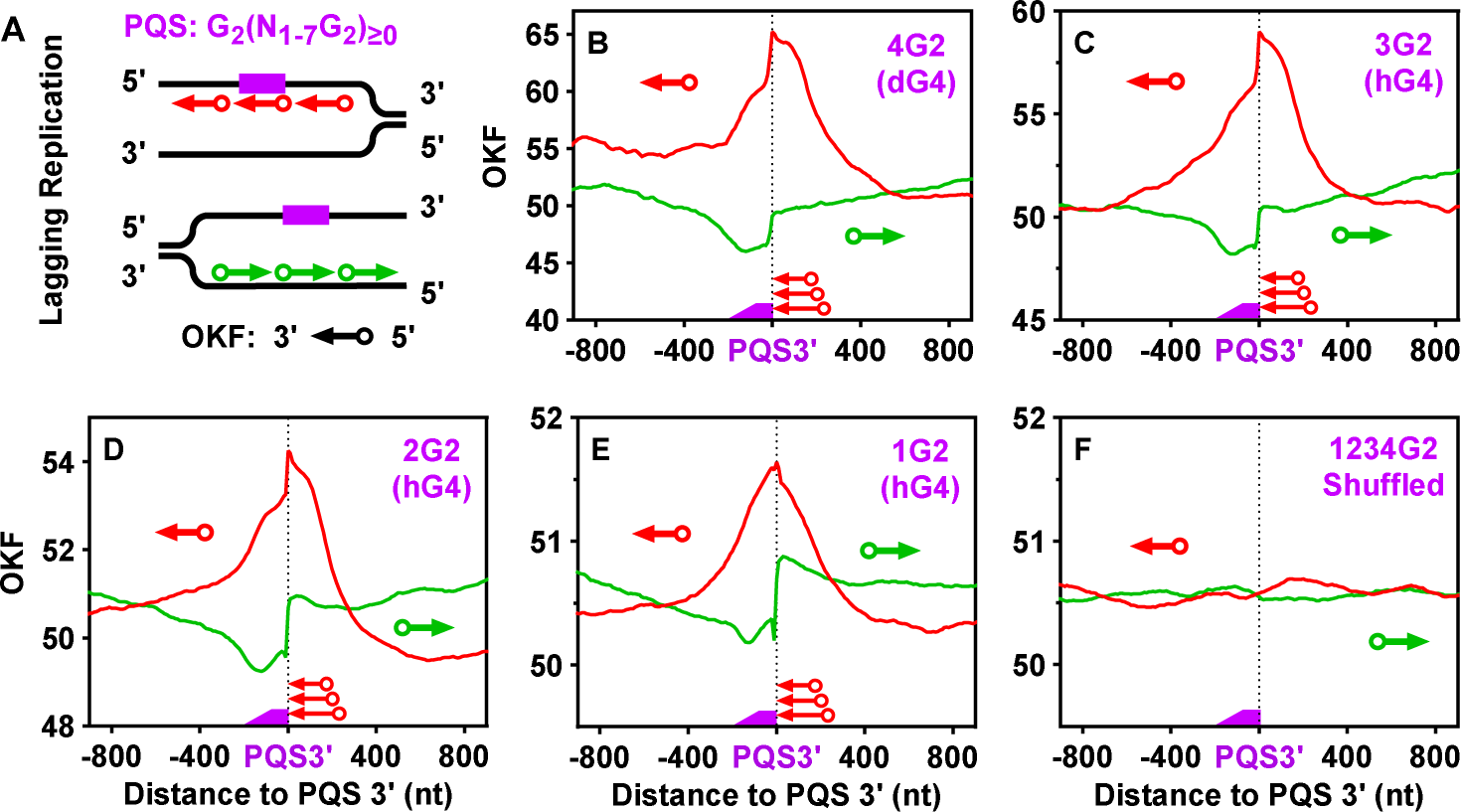
(A) Scheme of OKF distribution profiling on the PQS-bearing strand and the strand opposite to a PQS. (B-E) Distribution of OKFs at the 3’-end of PQSs with 1 to 4 or more GG tracts and capable of forming (B) dG4s or (C-E) hG4s of only two G-tetrads. (F) Distribution of the 3’-end of OKFs at the 3’-end of randomly shuffled PQSs.

**Figure 6.**
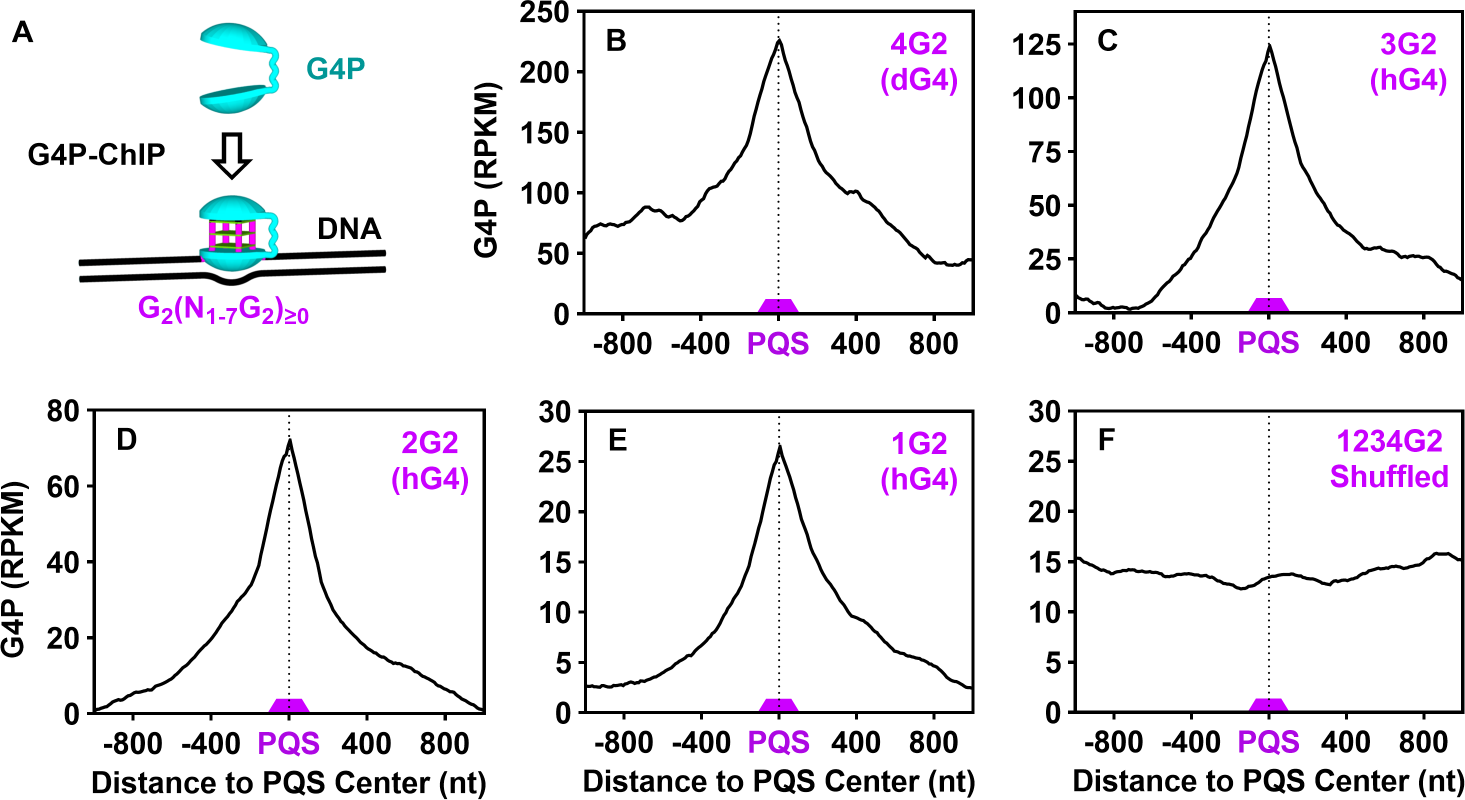
(A) Scheme of G4 detection by G4P ChIP-Seq. (B-E) Enrichment of G4P at PQSs with 1 to 4 or more GG tracts and capable of forming (B) dG4s or (C-E) hG4s of only two G-tetrads. (F) Distribution of G4P at randomly shuffled PQSs. PQSs are the same as in Figure 5.

### Presence of RNA G-tracts in hG4s

In the above results, the stalling of OKF synthesis next to the PQSs and the G4P enrichment at the PQSs (Figure 3-6, Figure S1-S4) provided clear evidence for the formation of hG4s at the PQSs with less than four G-tracts. Since these motifs alone were unable to form G4s, the involvement of RNA G-tracts is expected. To determine whether RNA was involved, we performed a two-step immunoprecipitation to sequence the RNA in the G4s (Figure 7A). A DNA antibody was used to capture cross-linked DNA fragments that could contain dG4s, hG4s, or R-loops (not shown in the scheme). The captured DNA was then further precipitated using a G4P-binding antibody or a non-specific antibody of the same origin as a control. The DNA in the precipitates was digested with DNase, and the remaining RNA was sequenced. As shown in Figure 7B, RNAs were detected and those with 1 to 3 G-tracts were enriched compared to the non-G4-forming C/A/T-rich controls. In addition, the degree of enrichment was positively correlated with the number of RNA G-tracts, which is consistent with the known fact that more G-tracts have a greater tendency to participate in hG4 formation (9). Similar result was obtained when RNA GG tracts and the corresponding control motifs were included (Figure S9). In this case, a lower enrichment was observed because G4s with shorter G-tracts are relatively much less stable (8,13).

**Figure 7.**
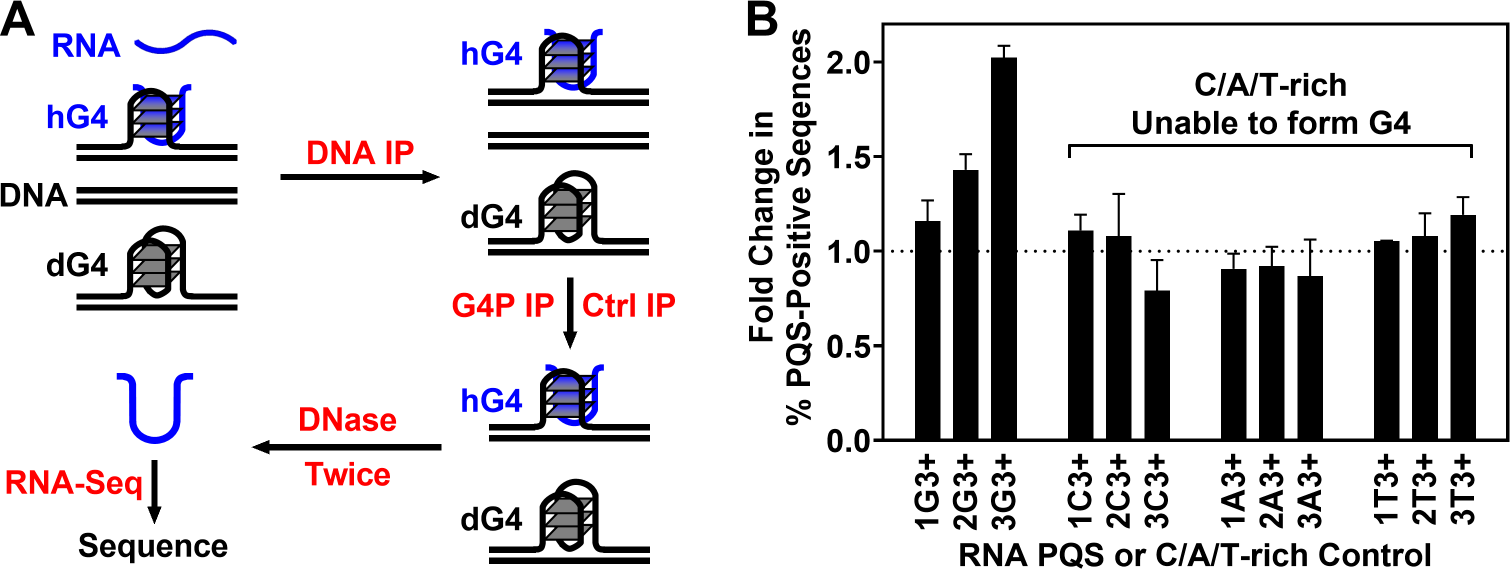
Detection of RNA G-tracts in hG4s. (A) Workflow of two-step immunoprecipitation (IP) and RNA sequencing. Precipitation of G4 with an anti-FLAG antibody (G4P IP) was performed in parallel with a non-specific antibody as control (Ctrl IP). (B) Change in percentage of G-tract-positive RNA sequences in G4P-bound DNA compared to control, expressed as mean ± SEM of three biological replicates.

### Interception of OKF synthesis by hG4s at PQSs

The OKFs are of different lengths, which was a source of variation that contributed to the OKF distribution profiles (Figure 3 and 5, Figure S1-S4). To determine the exact termination position of the OKFs, we extracted the coordinates of the 3’-end of the OKFs and plotted their distribution across the PQSs (Figure 8A and 9A, arrowhead). A sharp peak was observed at the 3’-end of the PQSs, regardless of whether the PQSs were able to form dG4 or hG4 with two (Figure 9, B-E) or more (Figure 8, B-E) G-tetrads. This sharp peak suggests that the syntheses of OKFs were intercepted at the 3’ side of the G4s in the PQSs. If no G4s were formed, then the 3’-end of these OKFs would terminate randomly instead. To validate this, we collected OKFs whose 3’-end was within a range of ±20 nts from the 3’-end of the PQSs and analyzed the location of both their 5’ and 3’-ends (Figure 10A). The results showed that the 5’ ends of these OKFs were distributed over a wide range of approximately 500 nts downstream of the PQSs (Figure 10, B-E). This feature indicates that the syntheses of these OKFs were initiated from a broad region and, according to their sharp 3’-end peak at the 3’-end of the PQSs, were abruptly interrupted by the G4s in the PQSs. On the other hand, the sharp peak of the OKF 3’-end with improved background (Figure 8 and 9) makes it a much clearer indicator of G4 formation.

**Figure 8.**
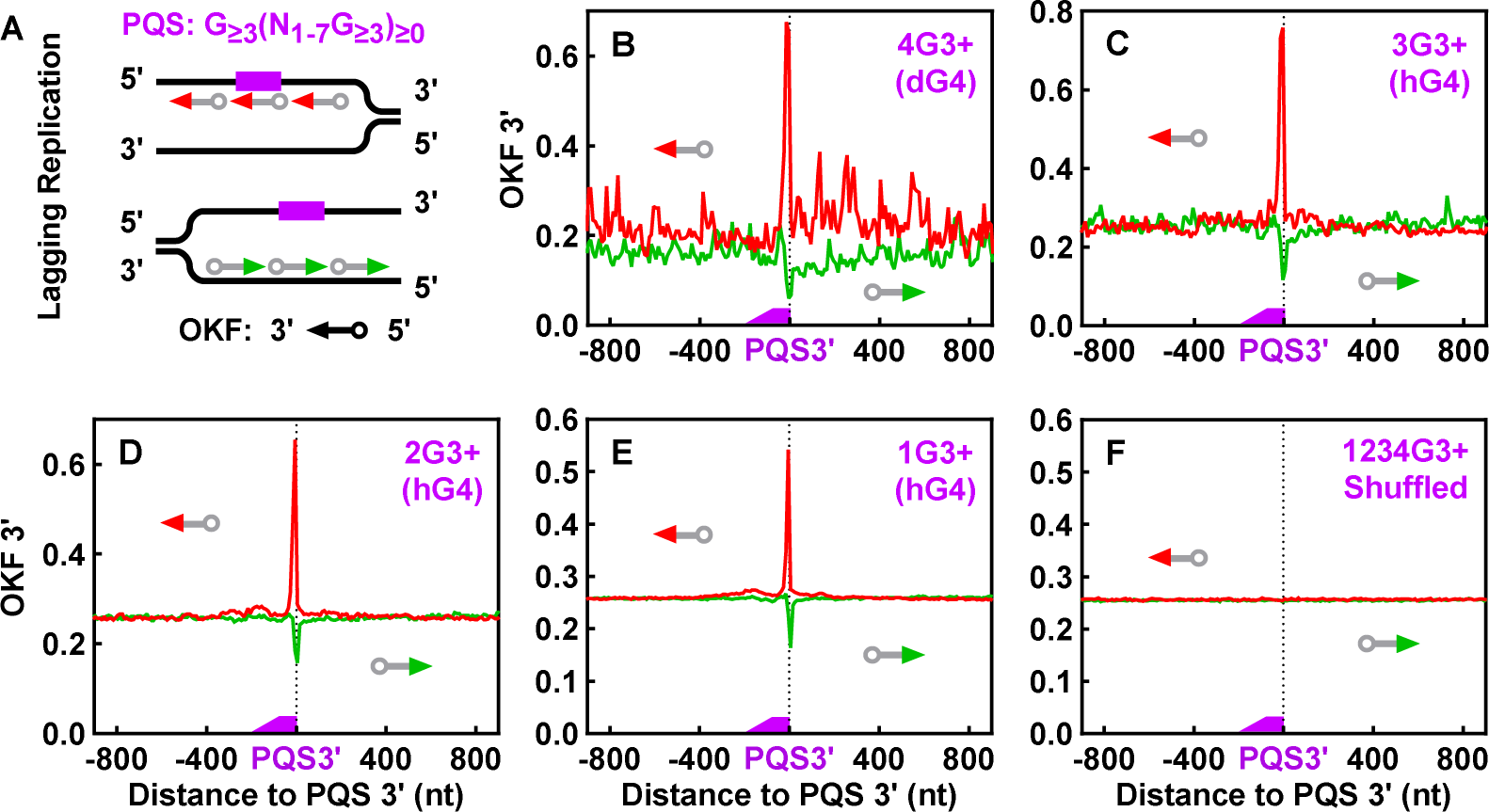
(A) Scheme of OKF 3’ distribution profiling on PQS-bearing strand and strand opposite to PQS. (B-E) Distribution of the 3’-end of OKFs at the 3’-end of PQSs with 1 to 4 or more G_≥3_ tracts and capable of forming (B) dG4s or (C-E) hG4s of three or more G-tetrads. (F) Distribution of the 3’-end of OKFs at the 3’-end of randomly shuffled PQSs. The PQSs are the same as in Figure 3. Note that the body and tail of the arrows are grayed out to indicate that only the 3’-end coordinate of the OKFs was considered.

**Figure 9.**
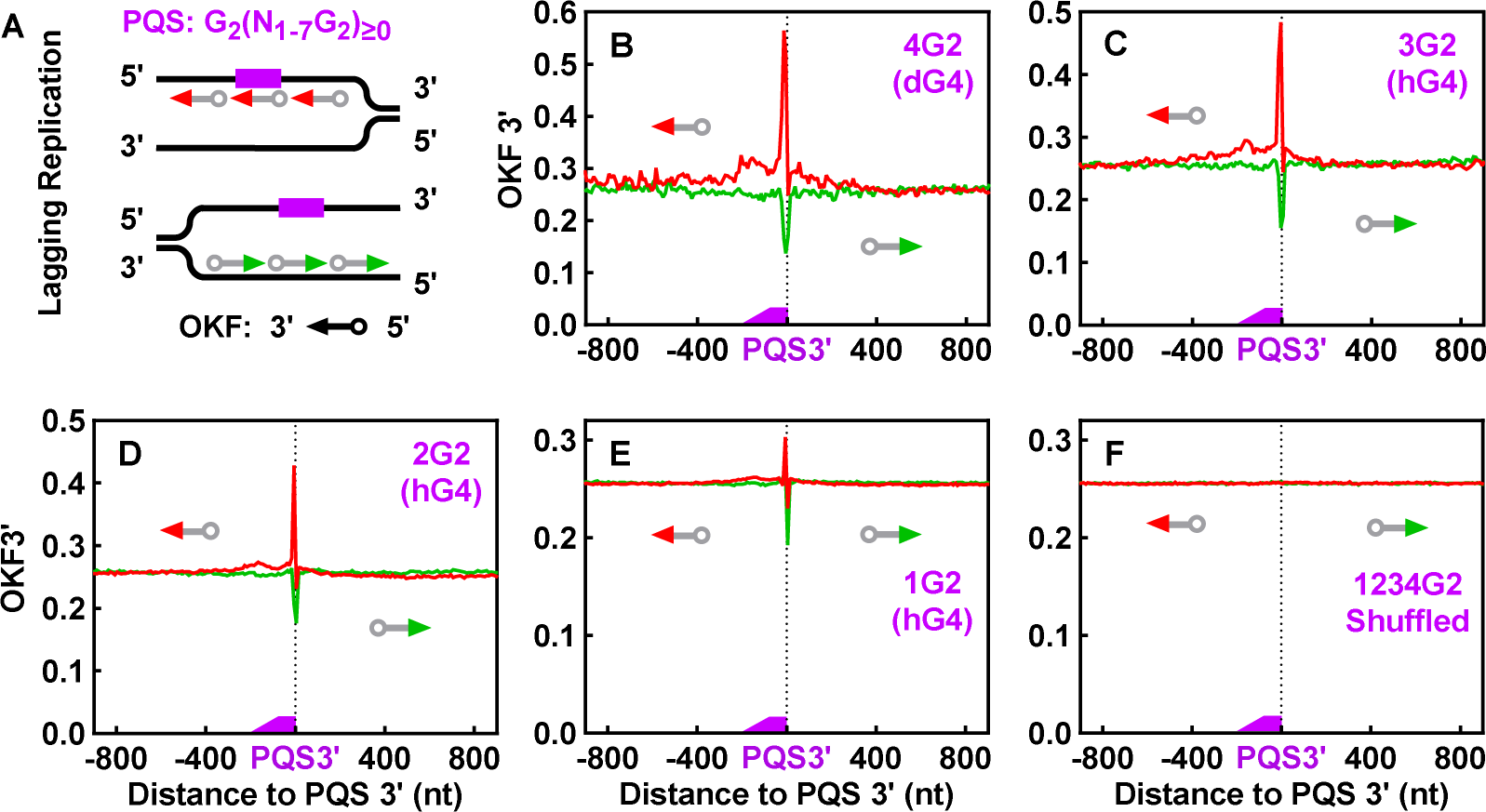
(A) Scheme of OKF 3’ distribution profiling on PQS-bearing strand and strand opposite to PQS. (B-E) Distribution of the 3’-end of OKFs at the 3’-end of PQSs with 1 to 4 or more GG tracts and capable of forming (B) dG4s or (C-E) hG4s of only two G-tetrads. (F) Distribution of the 3’-end of OKFs at the 3’-end of randomly shuffled PQSs. PQSs are the same as in Figure 5.

**Figure 10.**
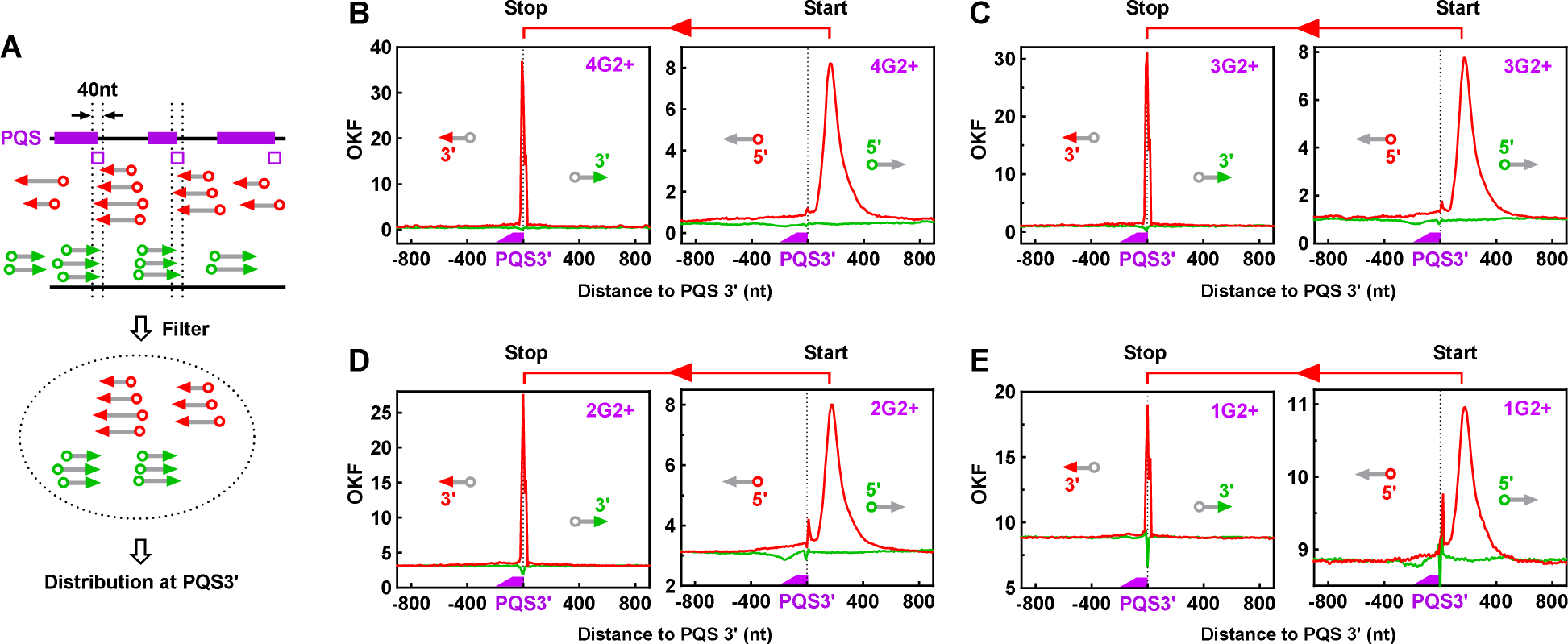
Interception of OKF syntheses at the 3’-end of PQSs. (A) Scheme of filtering of OKFs whose 3’-end was within the ±20 nts region (open box) of the PQS 3’-end. (B-E) Interception of OKF syntheses at the 3’-end of PQSs capable of forming (B) intramolecular dG4s or (C-E) hG4s of two or more G-tetrads. Bin size: 5 nts.

### The structural form of hG4s

In the polymerase stop assay, the number of stop sites is determined by the number of alternative G4s that can form with the available G-tracts (12) (Figure 2C). In the case of a single DNA G-tract, only one hG4 can form with three additional RNA G-tracts, so a single stop is expected. For a DNA PQS with four G-tracts, only one stop would occur if a single dG4 forms by consuming all four G-tracts (Figure 2A). However, when hG4s form, a maximum of four stops is expected, with each hG4 having one G-tract from DNA and three from RNA. This principle has allowed us to distinguish hG4 from dG4 in transcribed DNA both *in vitro* and in bacteria (12). Here, we used the same approach to further identify the composition of G4s based on the number of stop signals of OKF synthesis.

In Figure 11, we examined the extension of OKFs at single nt resolution, spanning an 80 nt region centered on PQSs containing 1-4 GG tracts, each with loops of identical size. For the PQSs with a single GG tract, a single prominent OKF 3’-end peak was observed, indicating the formation of a single hG4 consisting of one GG tract from DNA and three from RNA (Figure 11A). In Figure 11B, examples of different combinations of G-tracts from DNA and RNA for hG4 formation and the corresponding expected stops in OKF synthesis are listed. In Figure 11C, OKF syntheses across the PQSs with two GG tracts resulted in two major peaks that became more prominent when the loop size was increased from 1 to 7 nts for better resolution. This observation was consistent with the formation of two hG4s, each composed of one GG tract from DNA and three from RNA. The same rule extended to the PQSs with three GG tracts, for which three peaks were detected accordingly (Figure 11D).

**Figure 11.**
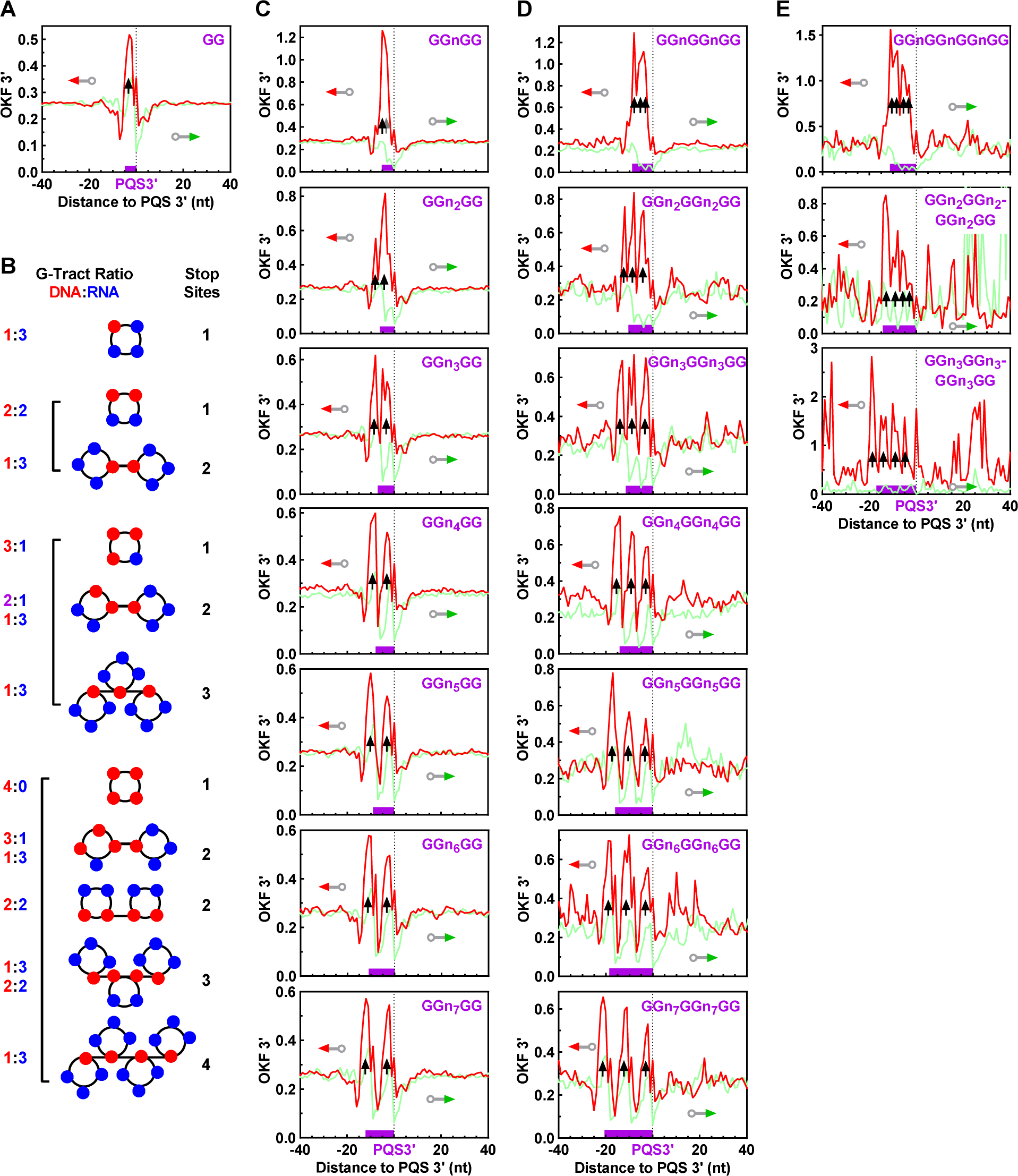
Distribution of OKF 3’-ends at PQSs with (A) one, (C) two, (D) three, or (E) four GG tracts showing the involvement of DNA G-tracts in hG4 formation. (B) Examples of combinations of DNA and RNA G-tracts in hG4 formation. “n” denotes any nucleotide except guanine. Bin size: 1 nt.

A PQS with four GG tracts would be expected to result in a single OKF 3’ peak if all GG tracts were used to form a single dG4. Interestingly, these PQSs instead showed four OKF 3’ peaks, indicating the formation of four hG4s (Figure 11E). This behavior was further confirmed in four additional datasets from independent studies (Figure 12), suggesting that a DNR:RNA G-tract combination of 1:3 was used to form each of the four hG4s (Figure 12, lower right scheme), although other minor forms of combinations could not be excluded. Similar results were obtained for the GGG (Figure S10) and GGGG (Figure S11) DNA tracts, all supporting the formation of hG4s.

**Figure 12.**
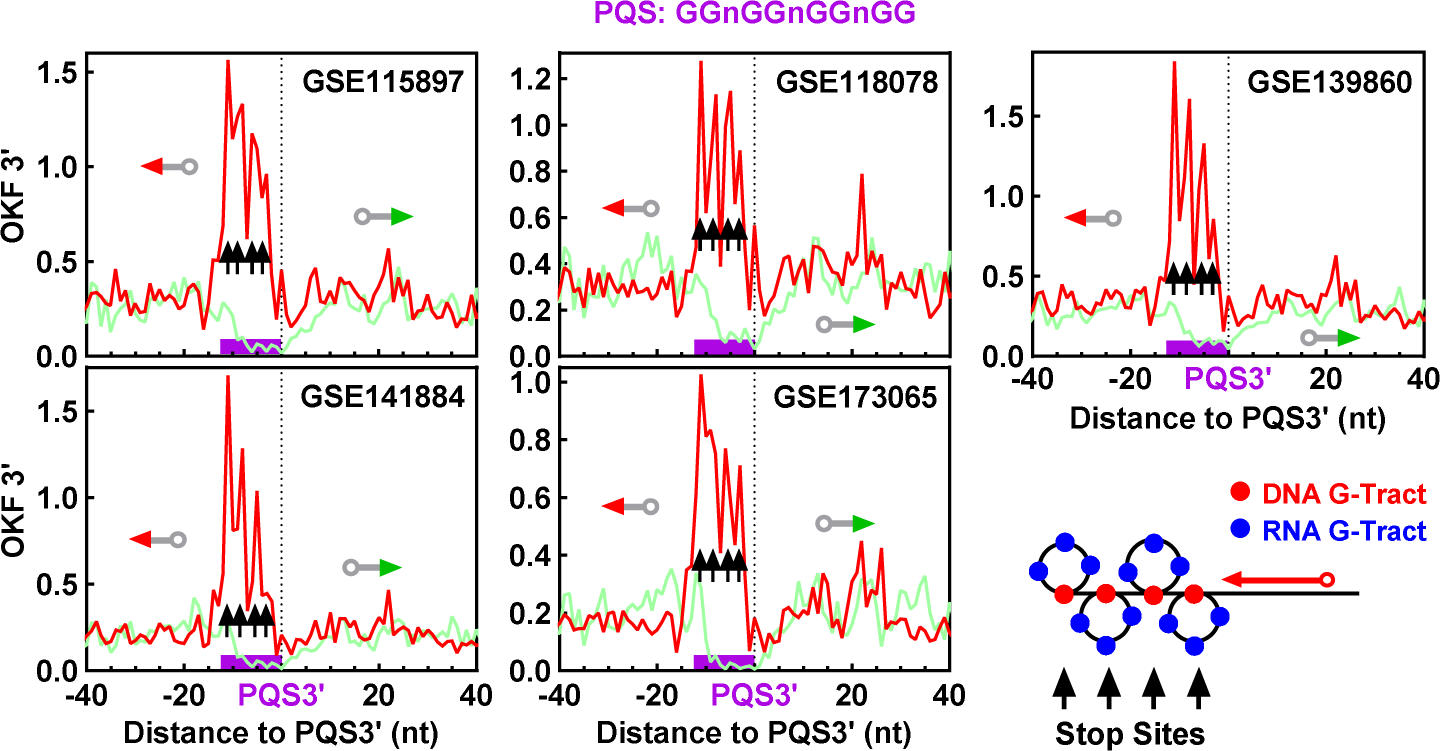
Distribution of OKF 3’-ends at PQSs with four GG tracts and 1-nt loops showing the involvement of DNA G-tracts in hG4 formation. The source of the original OK-Seq data is indicated in the panels. Bin size: 1 nt.

Taken together, our results in this section provide two important findings. First, even a single GG tract in the yeast genome can form an hG4. Second, a PQS with four G-tracts still forms hG4s, although it can simply form an intramolecular dG4 by itself. Judging from the well-resolved four dominating peaks (Figure 11E and Figure 12), the hG4s argued to be the major form of G4s. If not, a single prominent OKF 3’ peak would be expected at the first G-tract from the downstream side of the PQSs. A PQS in a chromosome is constrained in a more rigid DNA duplex, making the folding of G-tracts more challenging compared to a more flexible RNA strand. The dominance of hG4 formation in such cases was observed in our previous studies using linear duplex and plasmid DNA transcribed *in vitro* with T7 RNA polymerase (12,16), which was explained by the observation that hG4 formation is favored by faster kinetics and greater stability (16).

### Formation of hG4 in orphan PQSs

At this point, we were still concerned about the formation of hG4s in PQSs. A PQS is defined by a consensus that limits the loop between G-tracts to a maximum of seven nts, a rule that is generally accepted by the G4 community. However, when considering a PQS with fewer than four G-tracts, it is possible that neighboring G-tracts beyond the loop limit may be recruited to form a dG4 rather than an hG4. To address this concern, we identified three groups of PQSs carrying a single run of GG, GGG, and GGGG, respectively, separated from any other G_≥2_ tract by at least 20 to 100 nts. These G-tracts, termed “orphan G-tracts”, were analyzed for OKF stop and G4P binding signal (Figure 13).

**Figure 13.**
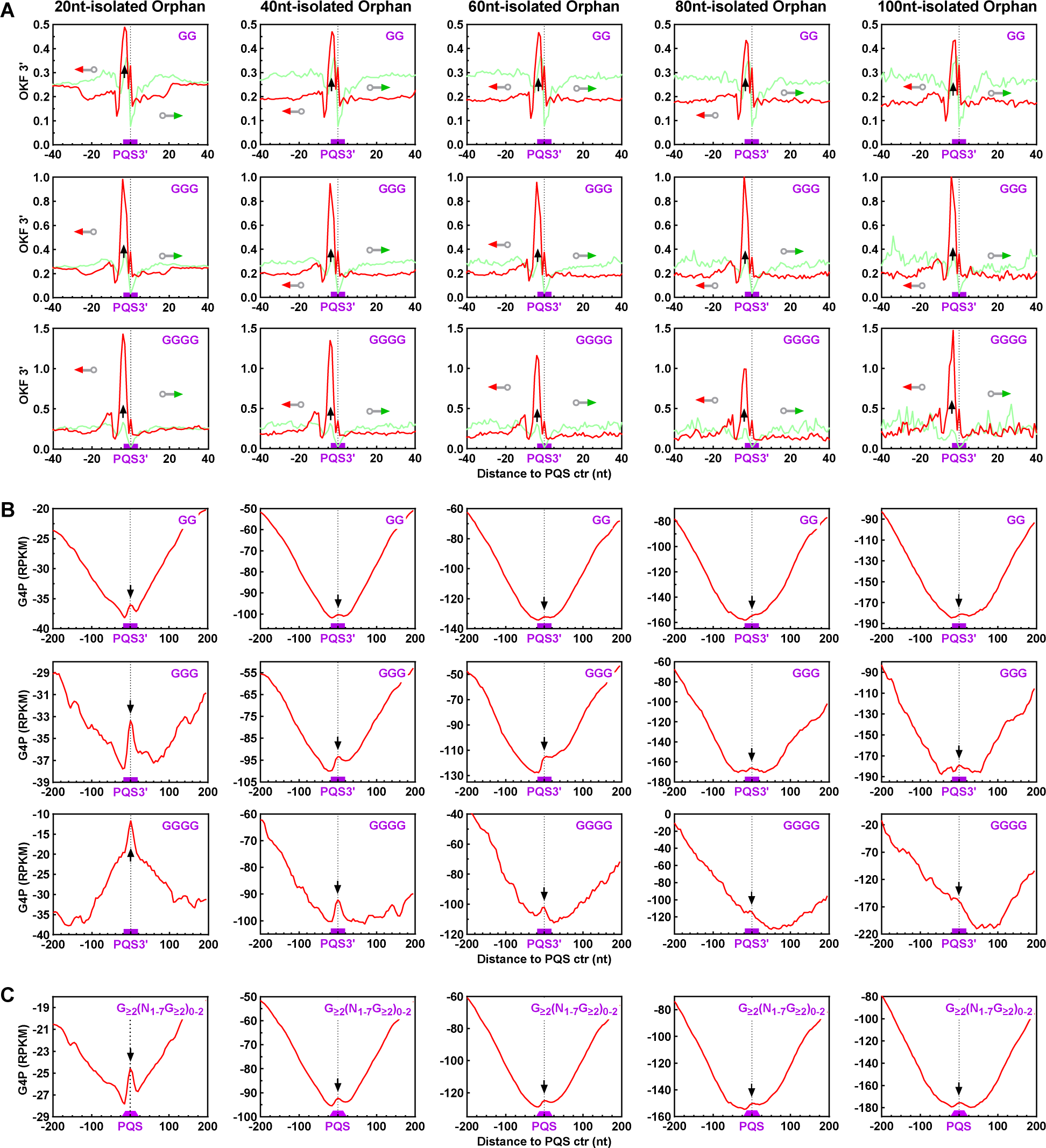
hG4 formation at orphan PQSs isolated from neighboring G-tracts by different numbers of nucleotides. (A) OKF 3’-end pausing at PQSs of single GG, GGG, GGGG tracts. (B) Detection of G4s by G4P at PQS of single GG, GGG, GGGG tracts. (C) Detection of G4s by G4P at PQSs with 1-3 G_≥2_ tracts. All PQSs are unable to form a G4 by themselves. Isolation sizes are given above the panels.

For all these G-tracts, a single prominent OKF 3’-end peak was detected as an indication of G4 formation. In particular, the peak height remained constant as the isolation flanks were extended from 20 to 100 nts (Figure 13A, panels from left to right), arguing against the possibility of dG4 formation. Otherwise, a decrease in G4 formation with increasing isolation size would be expected. In contrast, the peak height increased as the G-tract size increased from 2 to 4 nts (Figure 13A, panels from top to bottom), consistent with the well-documented fact that G4s with longer G-tracts are more stable (44). The formation of hG4s was further confirmed by the enrichment of G4P (Figure 13B), as indicated by the single G4P peak at most of the orphan G-tracts. The number of such orphan G-tracts rapidly decreased with increasing isolation size, leading to a degradation of the G4P signal. To improve the signal-to-noise ratio, we analyzed orphan PQSs with 1-3 G_≥2_ tracts that could not form a dG4 by themselves. In this case, a clearer G4P peak was detected for all five isolation sizes (Figure 13C). Taken together, these results from the orphan G-tracts/PQSs confirmed the ability of a PQS with less than four G-tracts to form an hG4.

### Survey of hG4 and dG4 sites in the yeast genome

Our analyses have revealed a distinctive picture that goes far beyond our previous perception of the nature and extent of G4 formation in the yeast genome. To get an overview, we surveyed the abundance of potential G4 sites in the genome and genes capable of forming either dG4s or hG4s with two or more G-tetrads. Since promoter activities are mostly concentrated in the 120 nts region upstream of TSSs in yeast (30,31), we merged this region with the gene body for analysis. According to the canonical consensus, the yeast genome can only form up to 38 dG4s with three G-tetrads. The discovery of hG4 formation increased the total number of G4 sites to 587,694, a >15,000-fold increase, of which canonical dG4s with two or more G-tetrads account for less than 2% (Figure 14A). Furthermore, hG4s are more abundant in genes, with a >50-fold higher abundance compared to dG4s (Figure 14, B versus C), highlighting the dominance and role of hG4s in transcriptional regulation.

**Figure 14.**
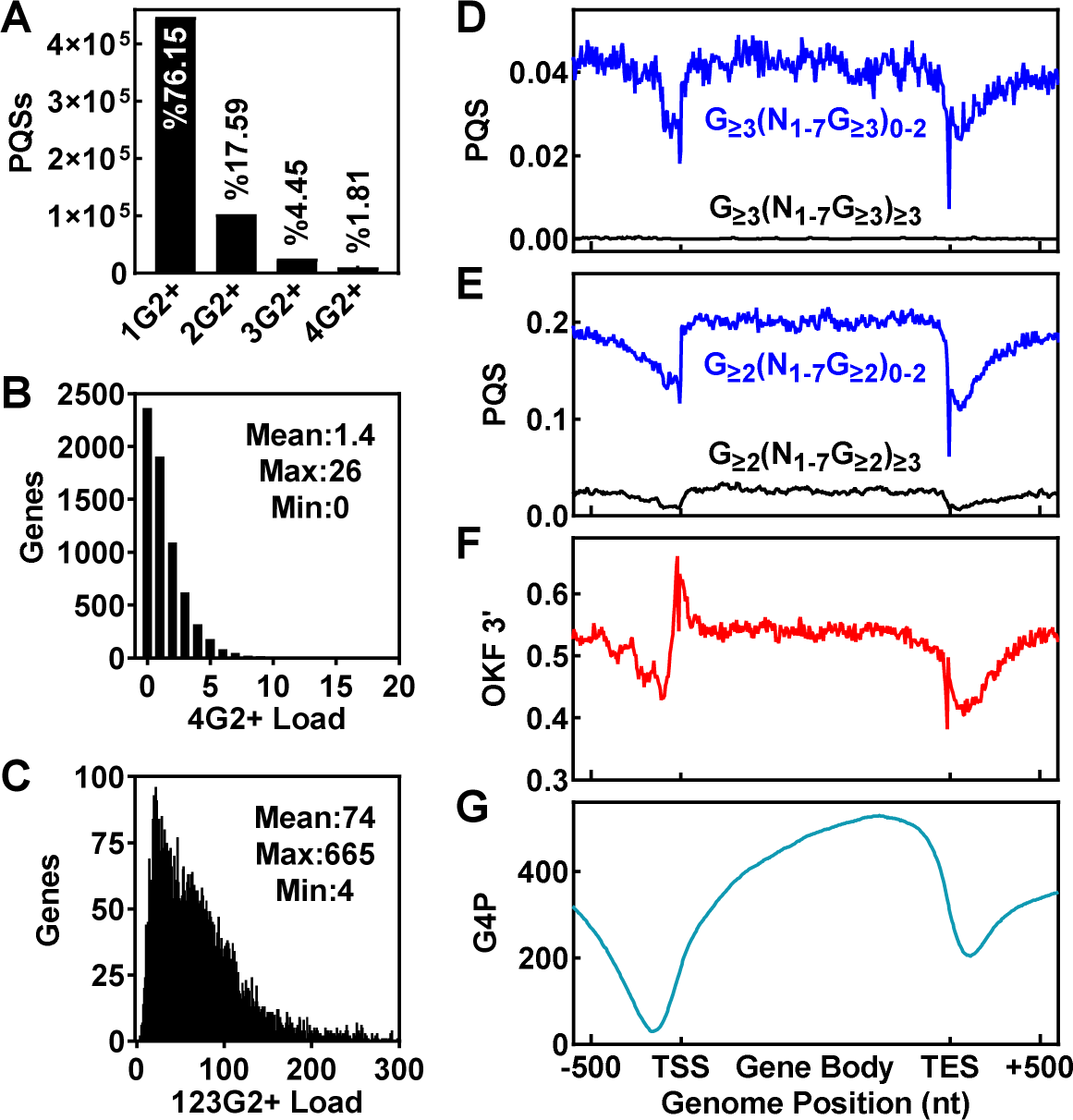
Overview of hG4 sites in the yeast genome and genes. (A) Counts of total PQS with different numbers of G-tracts. (B) PQS capable of forming dG4s in genes. (C) PQS capable of forming only hG4s in genes. (D) Distribution of PQS capable of forming only hG4s (blue line) and dG4s (black line) of three or more G-tetrads in genes. (E) Distribution of PQS capable of forming only hG4s (blue line) and dG4s (black line) of only two G-tetrads across genes. (F) Distribution of 3’-ends of OKFs across genes. (G) Binding of G4P across genes.

We also profiled the distribution of PQS, OKF, and G4P signals across genes. The results (Figure 14, D-E) showed that PQSs had a similar density throughout the genome, with a notable decrease near TSSs and transcription end sites (TESs). In contrast, the OKF 3’ signal showed its highest peak at TSSs (Figure 14F), probably due to higher production of RNA transcripts in this region. However, G4P binding instead showed a gradual decrease towards the TSSs (Figure 14G), which might be attributed to the increased activity of protein interaction and translocation associated with transcription initiation. The discrepancy between OKF 3’ and G4P in their distribution pattern suggests that OKF 3’ may be more sensitive and efficient than G4P in detecting genomic G4 formation.

## Discussion

In conclusion, we have successfully adapted an *in vivo* version of the polymerase stop assay with OK-Seq and used G4P to detect the formation of hG4s in the yeast genome, providing the first evidence for their existence and dominance in living eukaryotic cells. Together with our previous study demonstrating the formation of hG4s in bacterial cells, our work suggests that the dominant and genome-wide formation of DNA:RNA hG4s is a universal phenomenon in both the prokaryotic and eukaryotic kingdoms. The biological functions of G4s are diverse (23,45), and hG4s have several distinctive characteristics compared to canonical dG4s. First, hG4s are significantly more abundant because they can form with as few as one DNA G-tract. Second, hG4s also form and dominate in regions where DNA itself can form dG4s. Third, the different combinations of G-tracts of DNA and RNA G-tracts diversify the structural variability of hG4s (Figure 11B) in many ways (12). Most importantly, the tight coupling between hG4 formation and transcription (13,21) allows hG4s to mediate genome-wide transcription-dependent regulation.

G4s have been implicated in transcriptional regulation (46), with hG4s proposed to serve as constitutional cis-regulatory elements (9). Transcription-dependent hG4 formation may play a unique role in establishing a feedback loop for transcriptional regulation. G4s act as “roadblocks” that impede the translocation of proteins along a DNA strand (37,47), including RNA polymerase (9,12,48) and DNA polymerase, as demonstrated in this work. As a result, the presence of a G4 within a gene body results in transcriptional repression under both *in vitro* and *in vivo* conditions (9,12,49). An increase in transcription will lead to higher RNA levels and consequently higher hG4 formation, which in turn will lead to greater transcriptional repression. Given the widespread occurrence of hG4 formation in genes, this negative feedback may well serve as a common mechanism for maintaining transcriptional homeostasis and stability.

On the other hand, our study provides insight into the direct effect of hG4s on DNA replication. The stalling of DNA polymerase progression that we observed demonstrates how hG4s, acting as “roadblocks” on such a large genomic scale, can generally affect protein translocation along a DNA strand, a crucial event in genomic metabolism. Such effects may contribute to genome instability and explain why G4 stabilizing ligands cause DNA damage in a replication-dependent manner (50). In reality, an hG4 has the potential to affect any protein-DNA recognition or interaction, such as binding and diffusion.

In addition, the formation of hG4s of only two G-tetrads greatly diversifies the stability of G4 structures. Previous *in vitro* studies have shown that hG4s with two G-tetrads have a lifetime of minutes (8), while those with three G-tetrads can last for hours (13) if left undisturbed. Thus, these two sets of structures may play different roles, with the former being more responsive and the latter more persistent. Since the involvement of RNA G-tracts promotes G4 formation in both kinetics and stability (16), the combination of different ratios of DNA:RNA G-tracts in an hG4 should add another source of diversity.

Furthermore, the formation of hG4s may have played a role in evolution. PQSs capable of forming a dG4 are progressively selected during evolution, resulting in a higher frequency of these structures in higher species (3). From this perspective, the formation of hG4s in PQSs with 1-3 G-tracts of as few as two guanines could provide a smooth and necessary transition to facilitate the adaptation and selection of intramolecular dG4s. Without this gradual transition, the abrupt transition from nothing to the formation of canonical dG4s of four G-tracts would be difficult to justify.

Technically, the detection of G4s in living cells is challenging because G4s are dynamic structures that fold and unfold over time. Our work establishes OKF synthesis analysis as the only method that not only detects G4s, but also distinguishes their structural form in living cell genomes. G4 formation requires the opening of a DNA duplex, and our previous work has shown that G4 formation can be triggered by an approaching transcriptional bubble (35). A similar mechanism may also operate in a moving replication fork, where OKF synthesis and G4 formation occur simultaneously in close proximity, resulting in increased sensitivity. However, this mechanism may preferentially detect G4s in replicating DNA. It is important to note that transcription is an important driver of G4 formation (7). It is possible that G4s formed in regions that are actively transcribed but not replicated may be missed. Therefore, hG4s may form more extensively and abundantly in a genome than reflected by OKF synthesis.

## Acknowledgments

This work was supported by the National Natural Science Foundation of China (grant # 21977094 and 22037004) and the Shanxi “1331 Project”.

---

**Figure S1.**
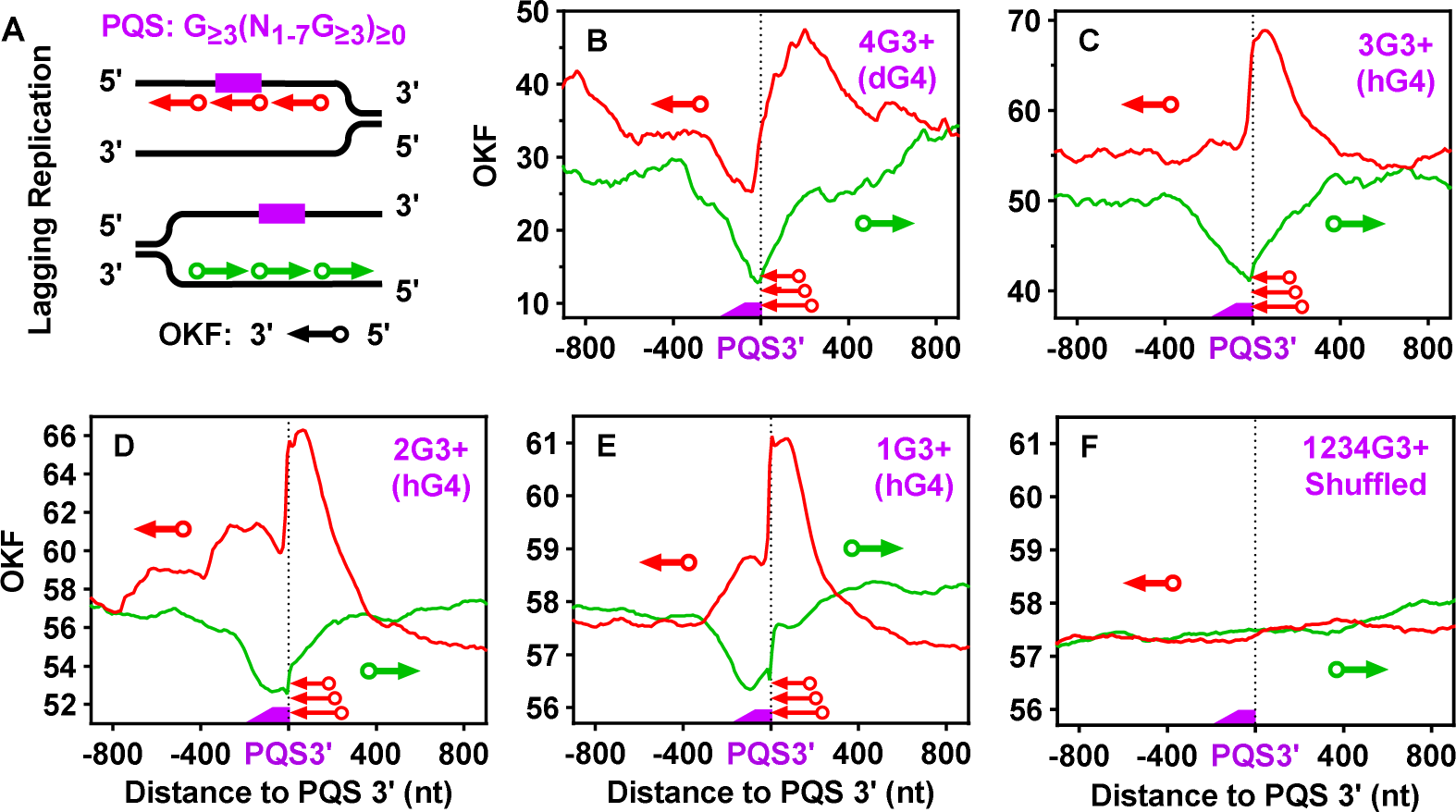
Distribution of OKFs at the 3’-end of PQSs with 1 to 4 or more G_≥3_ tracts that could form either dG4s or hG4s of three or more G-tetrads. Same as in Figure 3, except original OKF-seq data from GSE118078.

**Figure S2.**
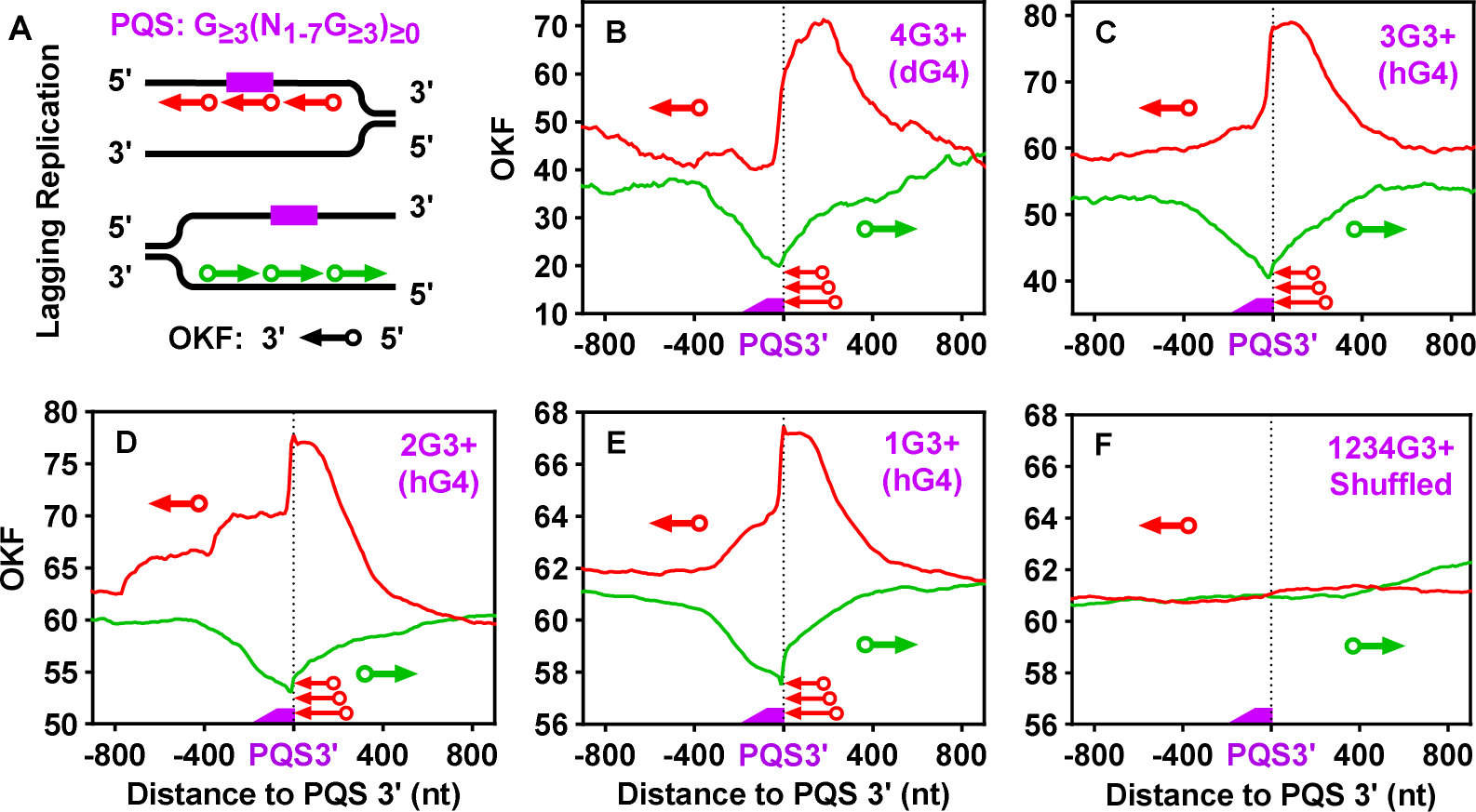
Distribution of OKFs at the 3’-end of PQSs with 1 to 4 or more G_≥3_ tracts that could form either dG4s or hG4s of three or more G-tetrads. Same as in Figure 3, except original OKF-seq data from GSE139860.

**Figure S3.**
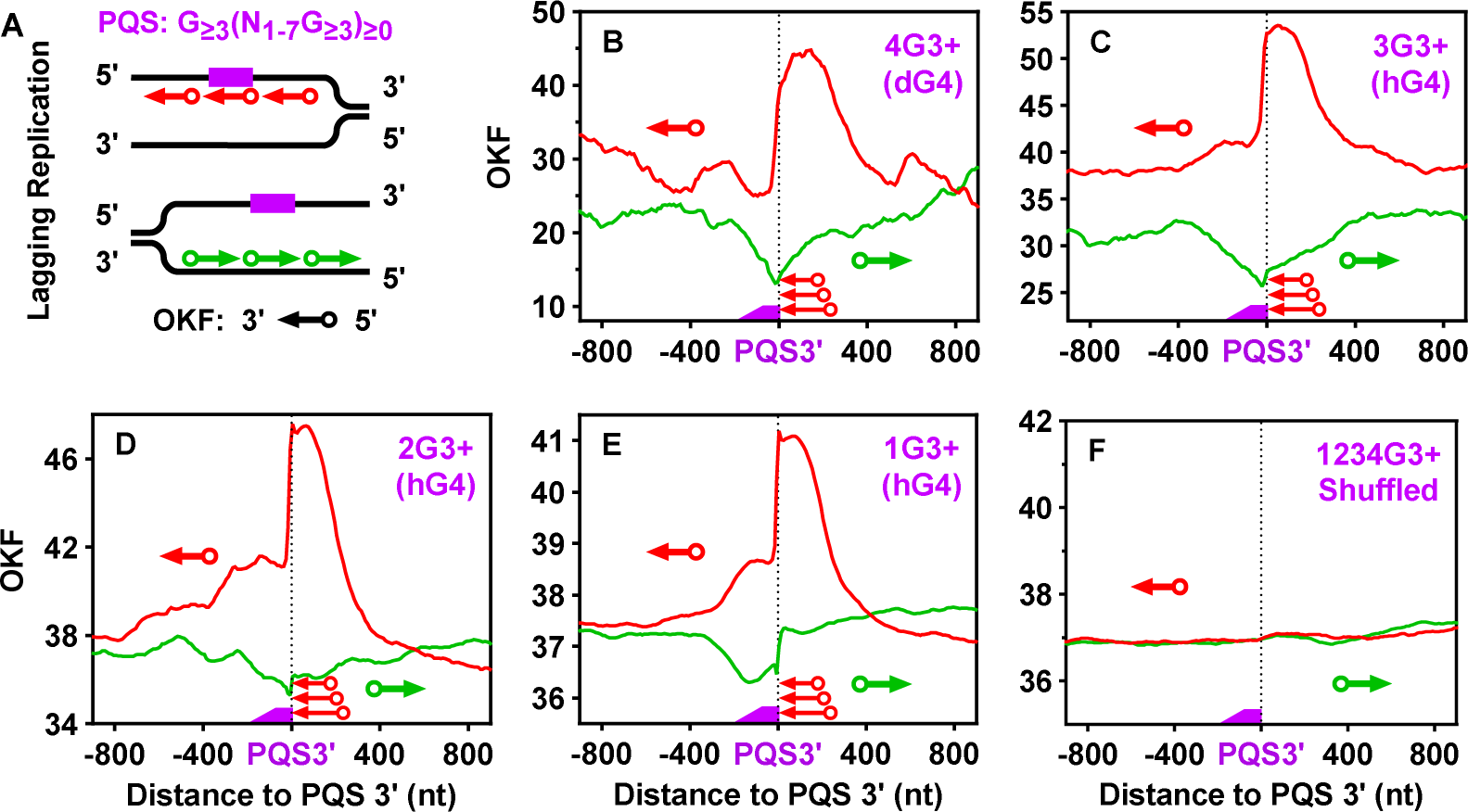
Distribution of OKFs at the 3’-end of PQSs with 1 to 4 or more G_≥3_ tracts that could form either dG4s or hG4s of three or more G-tetrads. Same as in Figure 3, except original OKF-seq data from GSE141884.

**Figure S4.**
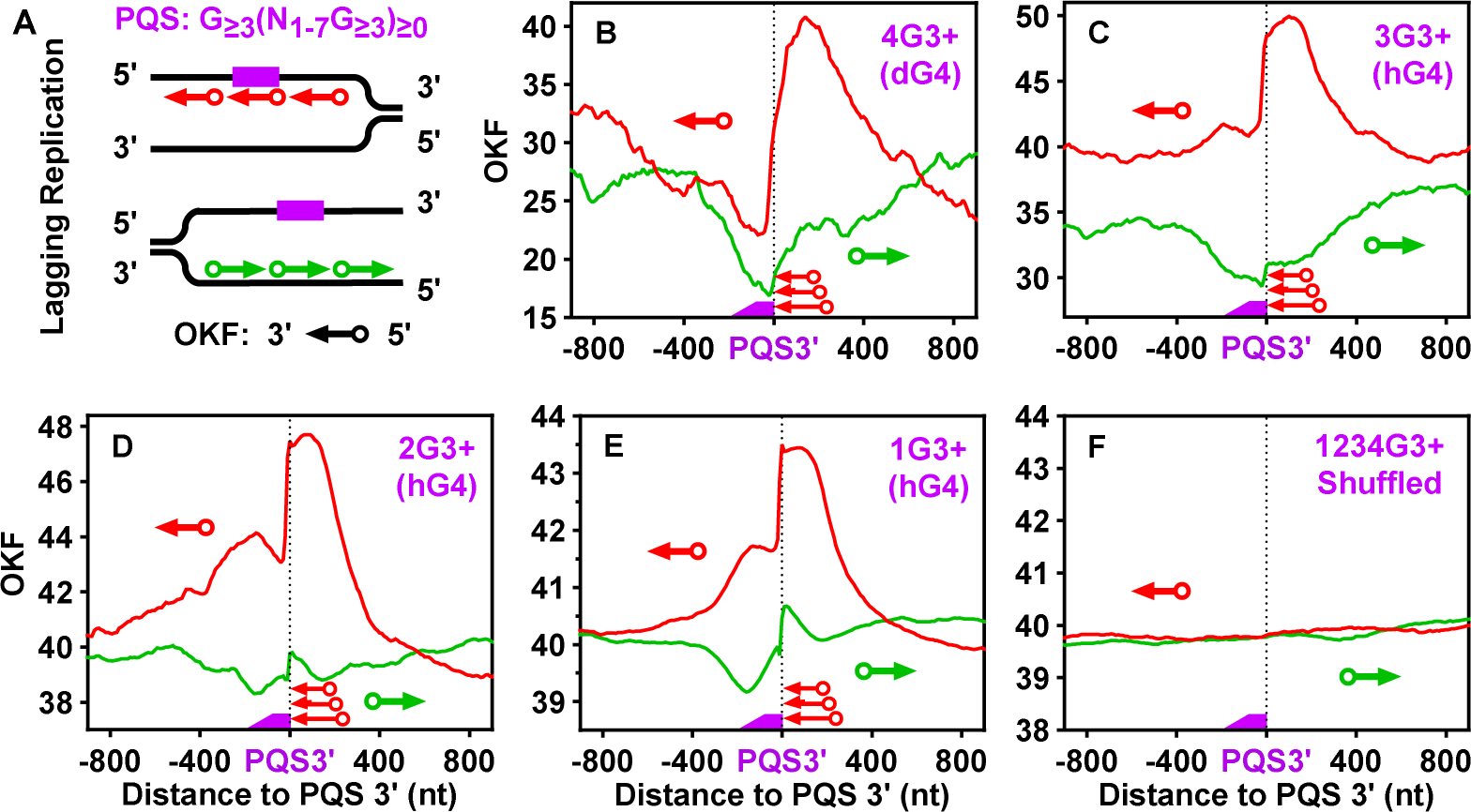
Distribution of OKFs at the 3’-end of PQSs with 1 to 4 or more G_≥3_ tracts which could form either dG4s or hG4s of three or more G-tetrads. Same as in Figure 3, except original OKF-seq data from GSE173065.

**Figure S5.**
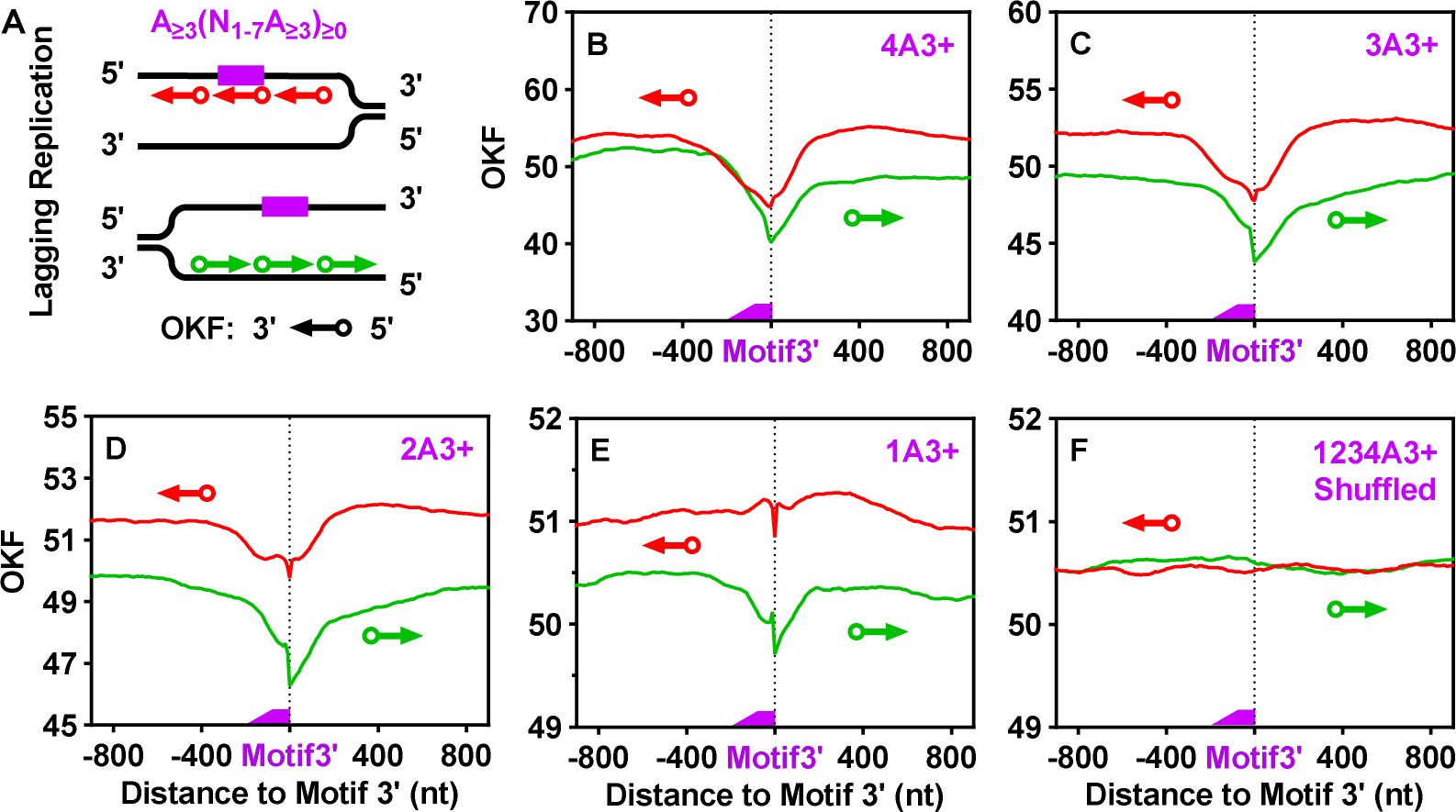
Distribution of OKFs at the 3’-end of A-rich motifs containing 1 to 4 or more A_≥3_ tracts that are unable to form G4. Same as in Figure 3, except that the PQSs were replaced by the indicated motifs.

**Figure S6.**
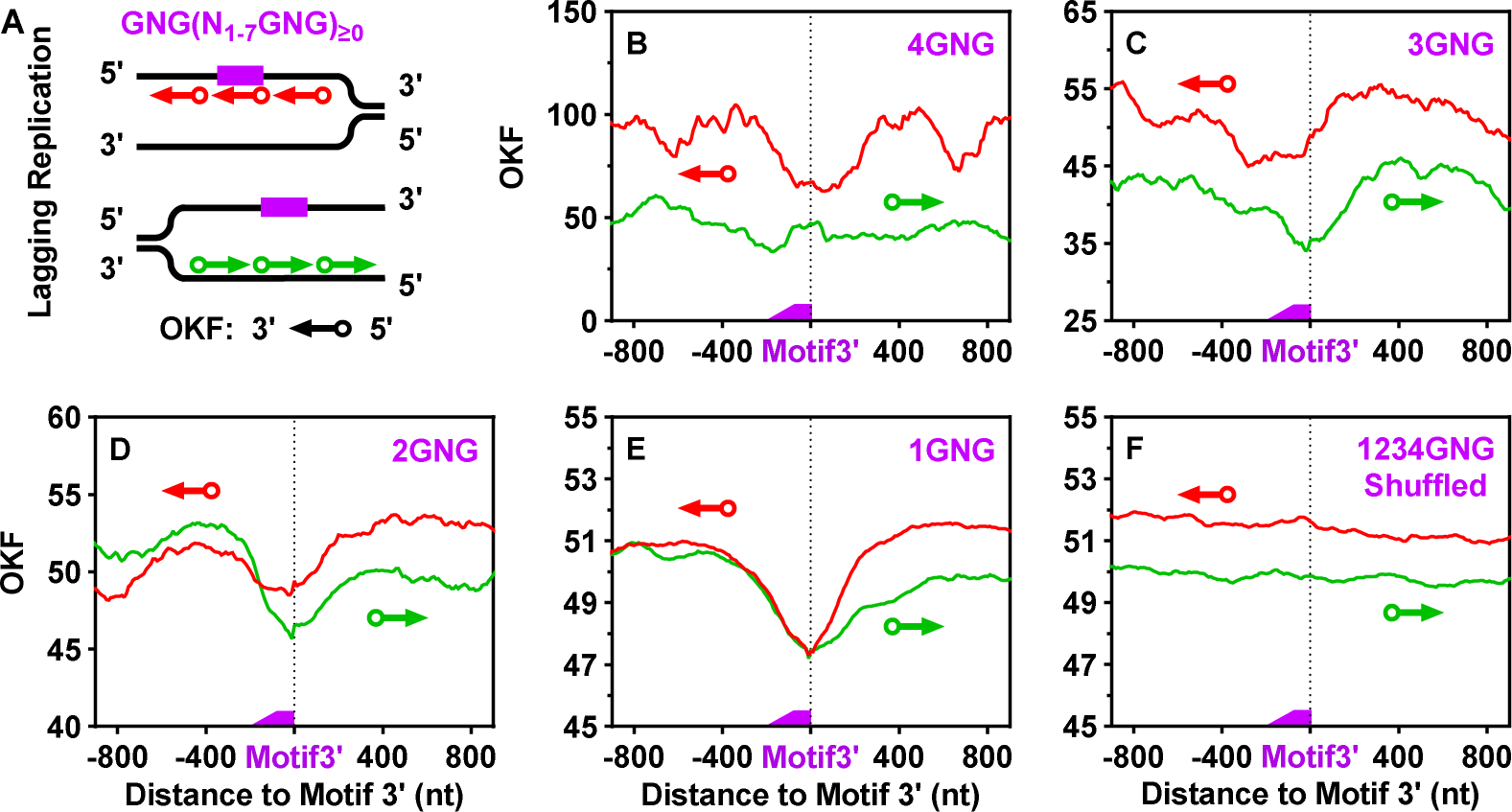
Distribution of OKFs at the 3’-end of motifs containing 1 to 4 or more GNG tracts that are unable to form G4. Same as in Figure 3, except that the PQSs were replaced by the indicated motifs.

**Figure S7.**
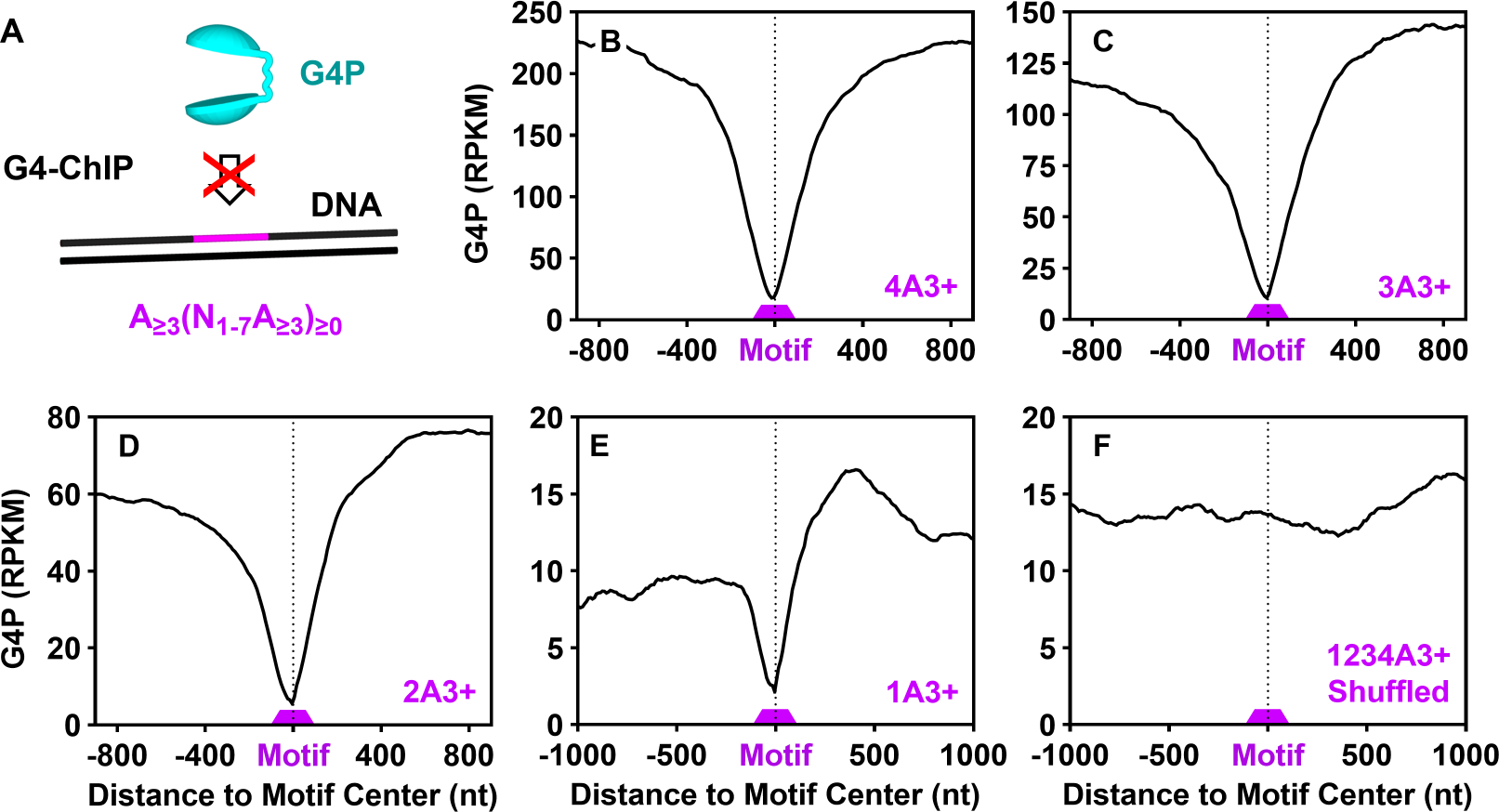
Binding profiles of G4P at A-rich motifs containing 1 to 4 or more A_≥3_ tracts that are unable to form G4. Same as in Figure 4, except that the PQSs were replaced by the indicated motifs.

**Figure S8.**
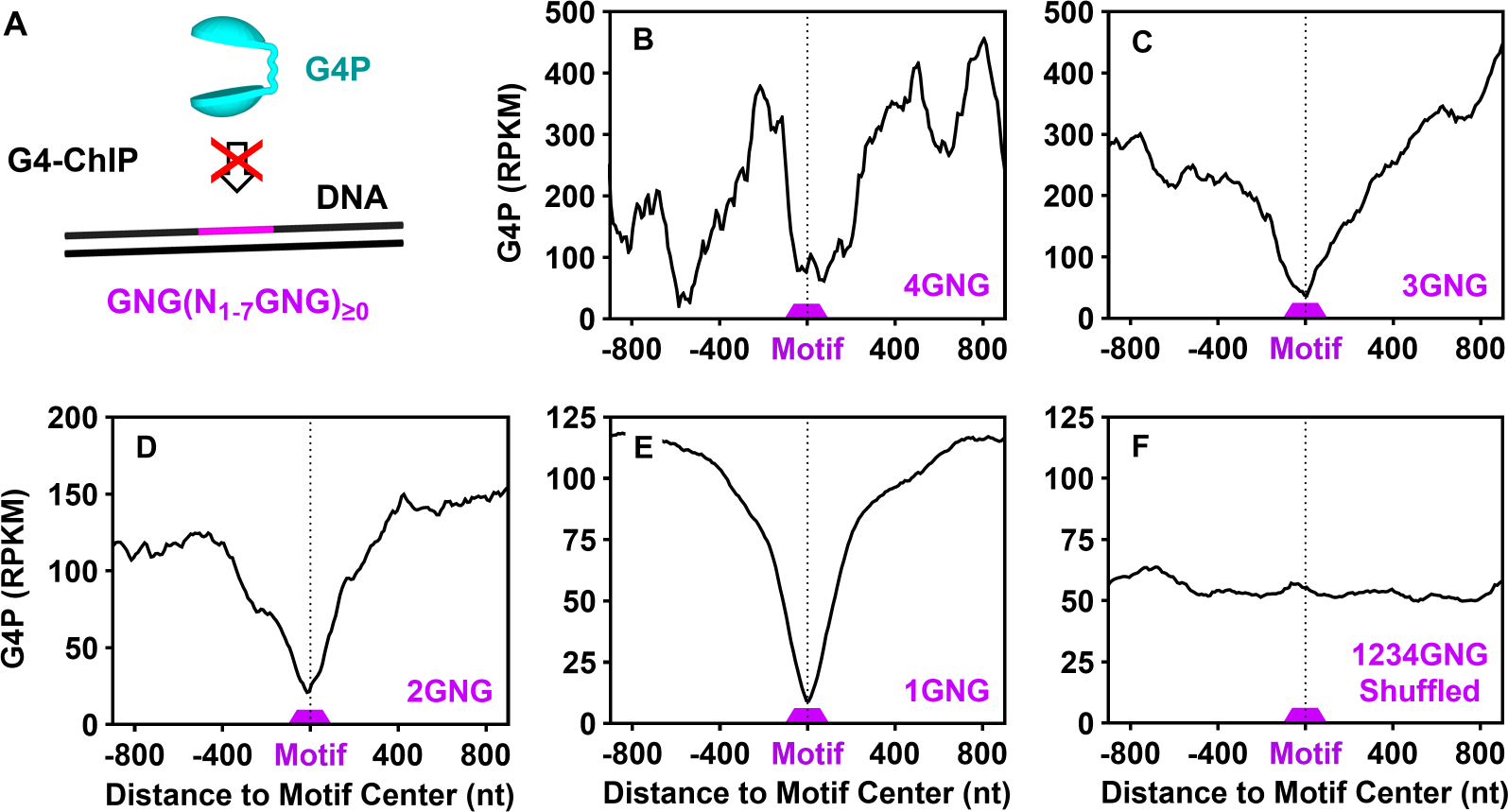
Binding profiles of G4P at motifs containing 1 to 4 or more GNG tracts that are unable to form G4. Same as in Figure 4, except that the PQSs were replaced by the indicated motifs.

**Figure S9.**
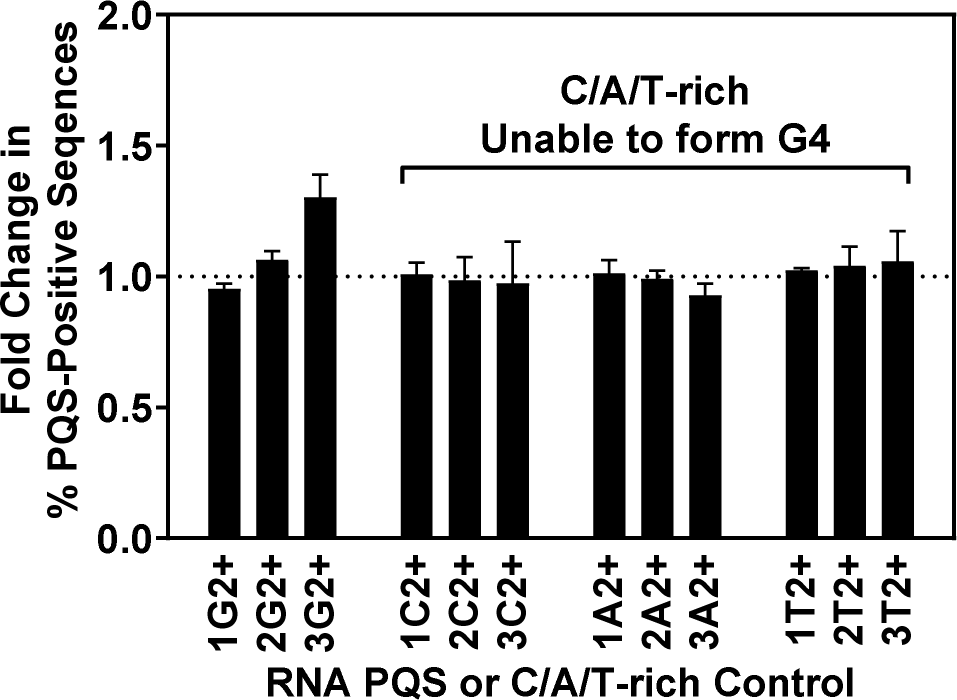
Detection of RNA G-tracts in hG4s with two or more G-tetrads. Same as in Figure 7B except for the size of the motif tracts.

**Figure S10.**
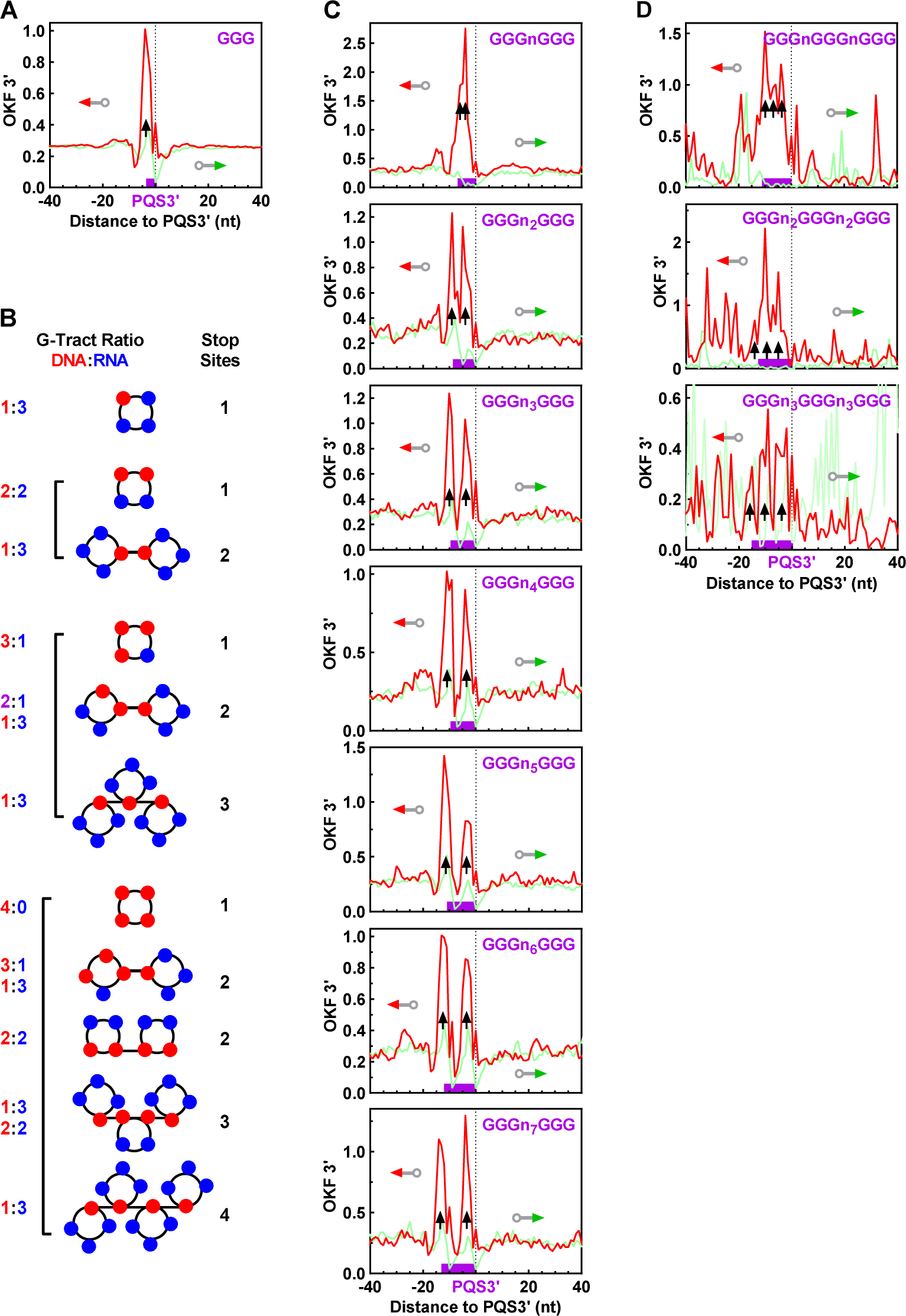
Distribution of OKF 3’-ends at PQSs with (A) one, (C) two, or (D) three GGG tracts. (B) Examples of combinations of DNA and RNA G-tracts in hG4 formation. “n” denotes any nucleotide except guanine. Bin size: 1 nt.

**Figure S11.**
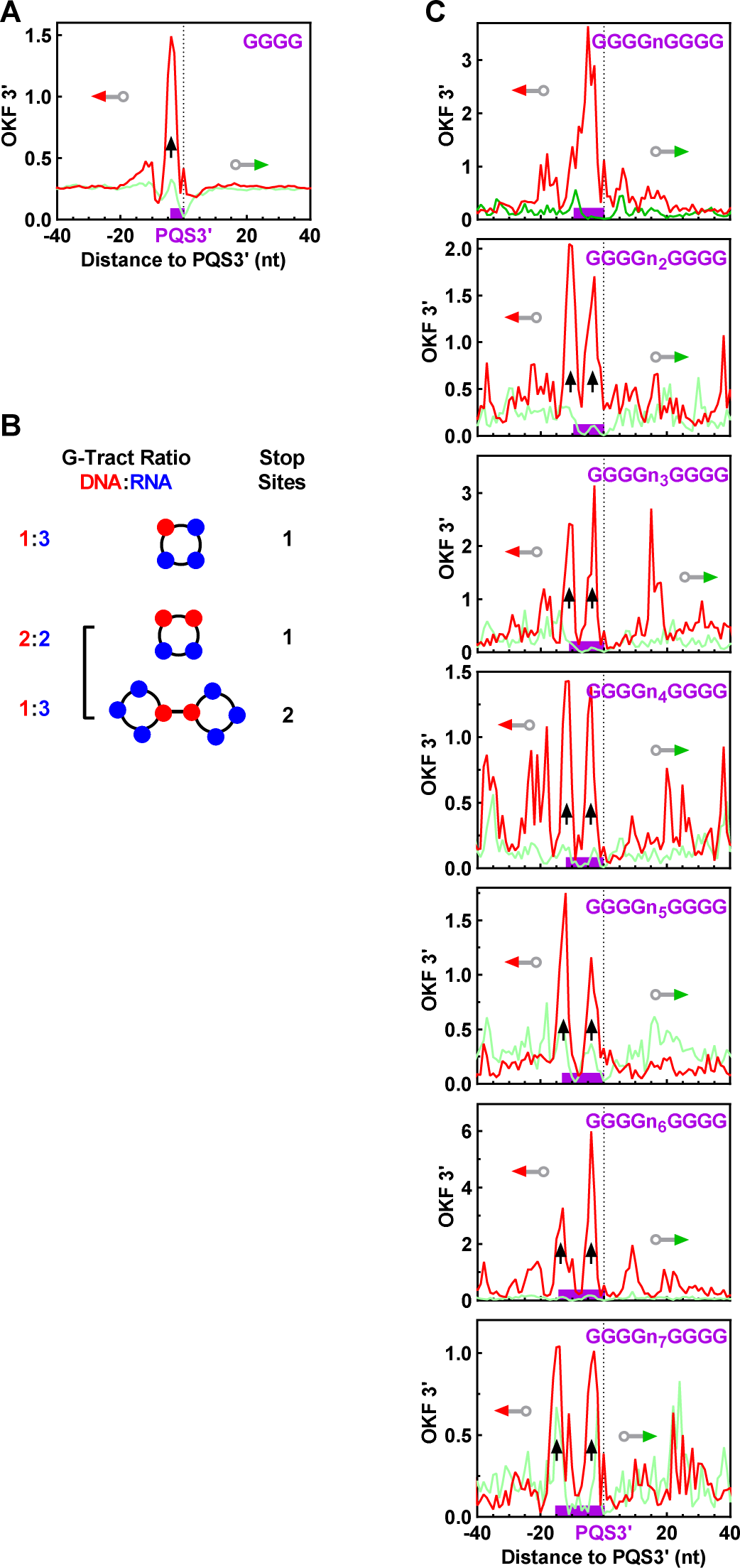
Distribution of OKF 3’-ends at PQSs with (A) one or (C) two GGGG tracts. (B) Examples of combinations of DNA and RNA G-tracts in hG4 formation. “n” denotes any nucleotide except guanine. Bin size: 1 nt.

**Table S1.**
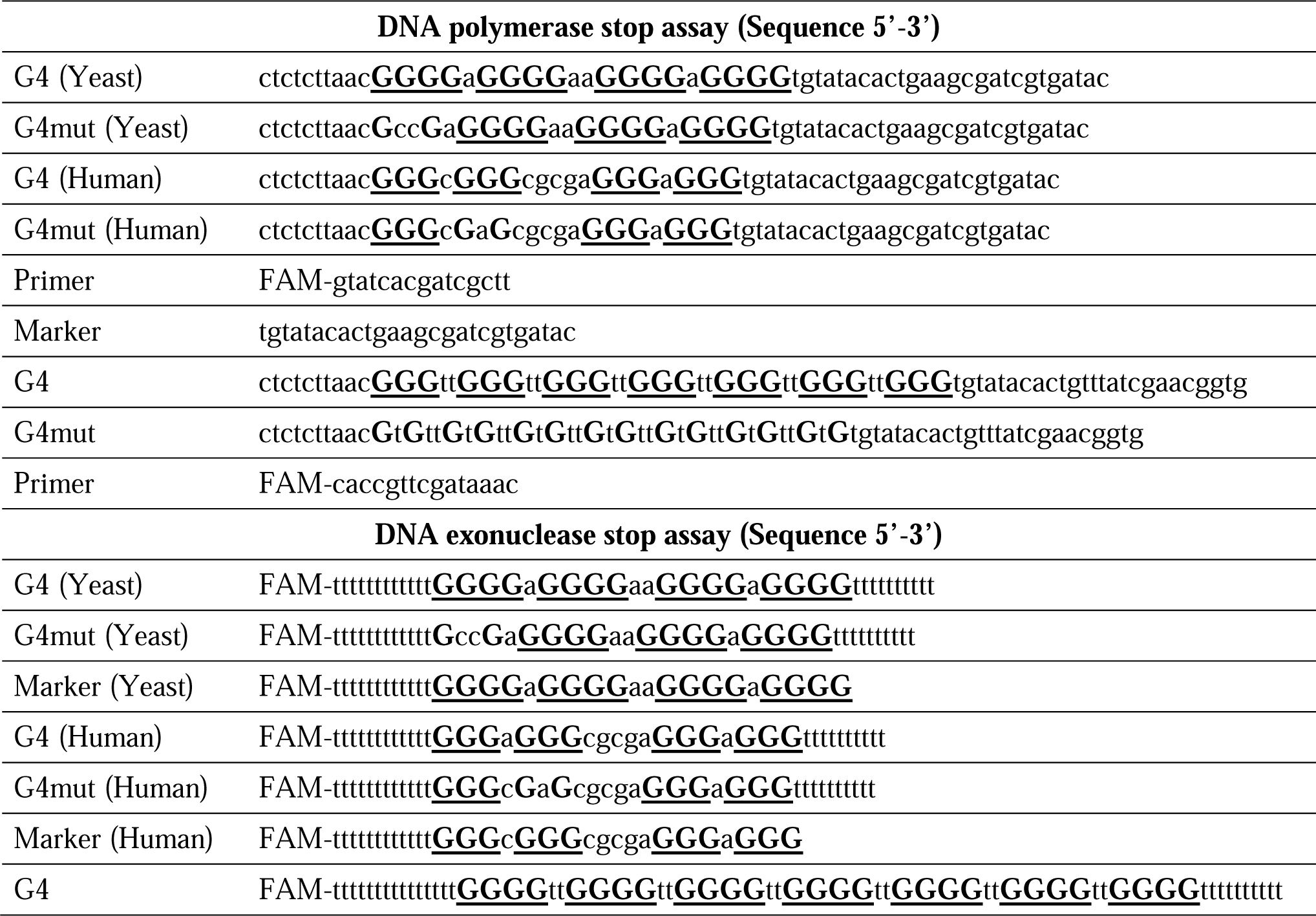
DNAs used in DNA polymerase as template and exonuclease stop assay as substrate.

## Notes

### Competing Interest Statement

The authors have declared no competing interest.

### Summary of Updates

Added new results on detection of RNA G-tracts in hG4s as Figure 7 and Figure S9; added new results as Figure 13C; added one author.

